# Spatially organized cellular communities shape functional tissue architecture in the pancreas

**DOI:** 10.1101/2025.04.16.649169

**Authors:** Alejo Torres-Cano, Jean-Francois Darrigrand, Gabriel Herrera-Oropeza, Georgina Goss, David Willnow, Anna Salowka, Debashish Chitnis, Morgane Rouault, Alessandra Vigilante, Francesca M. Spagnoli

## Abstract

Organ function depends on the precise spatial organization of cells across multiple scales, from individual cells to cellular communities that form specialized local niches and, ultimately, complex higher-order structures. While the identities of individual cell are increasingly well-defined, our understanding of how these diverse cell types are spatially distributed and communicate remains incomplete. In this study, we combine single-cell and spatial transcriptomic analyses to map pancreatic cell populations across space and time, from embryonic development to adult homeostasis in mice. Using these comprehensive maps, we systematically resolve spatial heterogeneity among pancreatic cell types and uncover basic tissue niches, which emerge as epithelial-mesenchymal units. We further characterize these niches functionally in both mouse and human models. We demonstrate that the mesenchymal lineage initially diversifies into various subtypes with specialized supportive roles during embryonic development. However, this complexity gradually diminishes over time, ultimately converging into a limited number of fibroblast sub-types in adult tissue. Our findings shed light on how different progenitor lineages co-develop and organize into structured communities that establish a mature, functional pancreas. This foundational framework could inform strategies for *in vitro* organogenesis and tissue-engineering in the context of pancreatic diseases.

## Main Text

Organs are composed of multiple cell types that develop in concert and organize into distinct tissue architectures, following a hierarchical spatial arrangement that is essential for their specialized functions(*1, 2*). Recent systematic single-cell RNA sequencing (scRNAseq) efforts have provided in-depth insights into the cellular composition of most embryonic and adult tissues(*3, 4*). However, the lack of spatial context in these analyses limits our understanding of how individual cells interact and arrange themselves to form and maintain functional tissue architectures. To address these questions, we need to shift our focus from individual cells to cellular communities on a collective scale. Functional tissue units (*a.k.a.* niches) arise from the dynamic interactions and coordinated actions of all constituent cells, requiring a holistic approach that encompasses the entire tissue topology(*5*). Furthermore, resolving the spatial organization of different progenitor cells and their microenvironment, as well as understanding how this organization changes during development, will provide insights into the mechanisms that regulate cell fate specification and differentiation. This will ultimately help linking cellular position to the emergence of distinct cell identities.

Here, we focus on the pancreas to interrogate the multicellular-scale mechanisms that define functional tissue architectures during organ development. The pancreas is an organ with both exocrine and endocrine functions, which are fulfilled by different cell types organized in distinct arrangements and locations across the tissue(*6*). Both exocrine and endocrine units arise from one common pool of endodermal progenitors, which progressively undergo differentiation and arrange into their respective structural and functional organization, either duct networks connected to acini or endocrine clusters (*a.k.a.* islets)(*6, 7*). These spatial arrangements are driven by multicellular interactions among pancreatic epithelial cells and the surrounding microenvironment, which is composed of a mix of different cell types, including mesenchymal, endothelial, immune cells and a rich extracellular matrix (ECM)(*8–11*). To date, the hierarchy and rules of interactions between the different cellular components that eventually result in emergent properties of tissue structure and function as pancreas development proceeds are yet to be defined. We hypothesize that an accurate cartography of the cell types populating the embryonic pancreas would illuminate uncharacterized interactions, as well as determine the modules responsible for establishing and maintaining its functional architectures.

### High-resolution map of the mouse embryonic pancreas

To visualize the distribution of the different cell types that constitute the embryonic pancreas and define the cellular interactions underlying its spatial organization, we generated a high-resolution transcriptomic map using direct RNA hybridization-based *in situ* sequencing (dRNA HybISS, Cartana part of 10X Genomics)(*12*), which enables multiplexed transcript detection at single-cell resolution (Fig. 1A). Specifically, we applied the HybISS technology to mouse pancreatic tissue at three embryonic stages (E) 12.5, E14.5 and E17.5 (Fig. 1A). Dorsal and ventral embryonic pancreata were collected from multiple independent embryos and cryosectioned at different positions along the anterior-posterior (AP) axis. We generated two custom panels of probes to map the expression of genes associated with microenvironmental cell types based on clustering annotation in our scRNAseq datasets, together with known pancreatic cell type markers (Fig. 1A-B; fig. S1) (table S1).

**Fig. 1.**
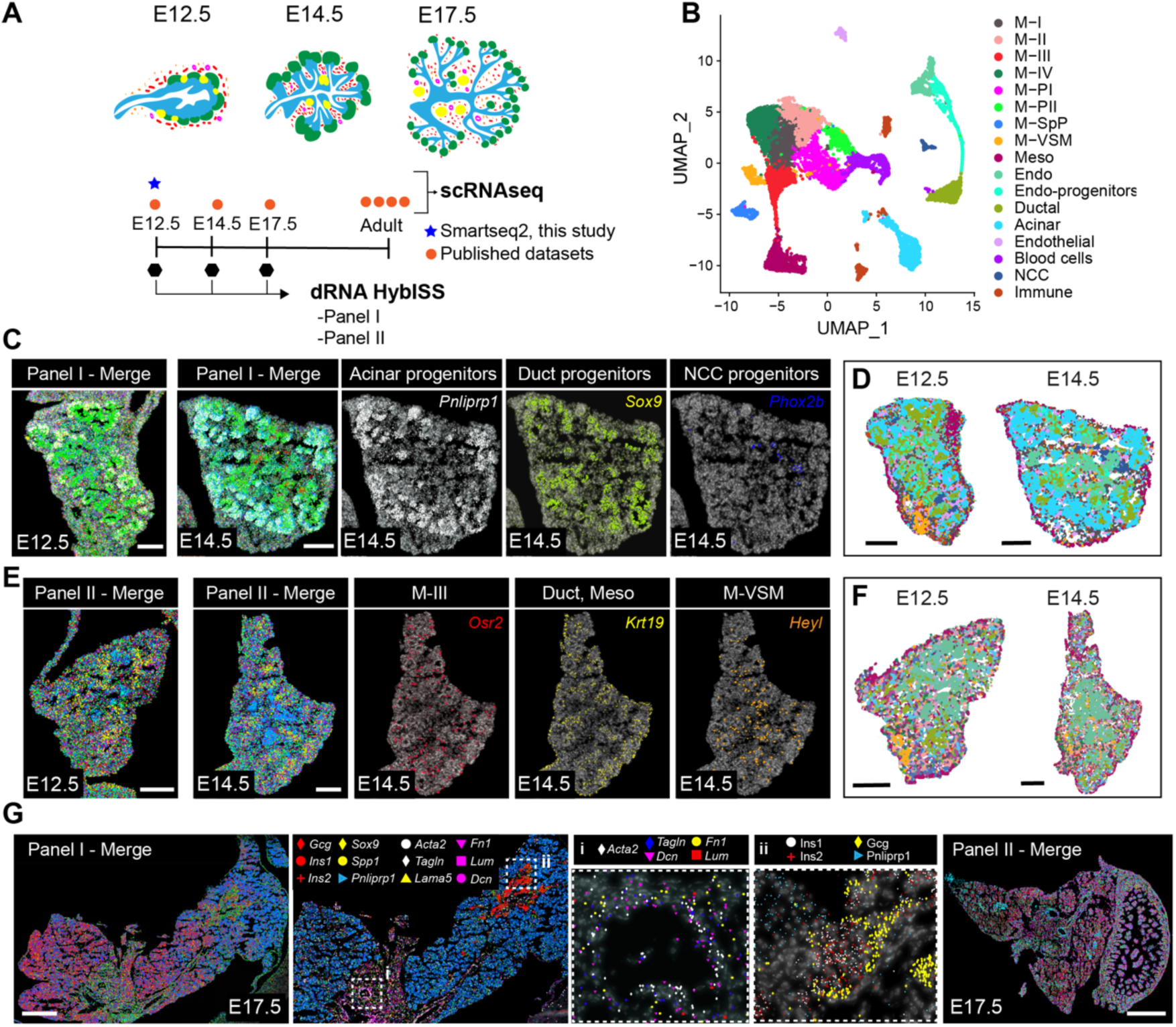
Major cell types and their spatial organization in the mouse embryonic pancreas as revealed by Hybridization-based in situ sequencing (HybISS). (**A**) Overview of the experimental design and sample collection strategy. HybISS experiments were performed using two gene panels (Panel I and II) targeting distinct sets of cell types in the embryonic pancreas. sc-RNAseq datasets from embryonic pancreatic tissue at the indicated stages and adult tissue were used for gene panel selection and subsequent analysis. (**B**) Uniform Manifold Approximation and Projection (UMAP) plot showing clustering of sc-RNAseq profiles of embryonic pancreatic tissue at E12.5, E14.5 and E17.5. Publicly available sc-RNAseq datasets(*16*) were integrated with high-coverage Smart-seq2 that we performed on FACS-isolated GFP^+^ cells from E12.5 pancreas of transgenic (Tg) Prox1-GFP, *Nkx2.5*-Cre;*R26*mTmG and *Nkx3.2*-Cre;*R26*mTmG Tg embryos (see fig. S1). Clusters are colour coded to indicate their annotated cell type. (**C**) Representative HybISS images showing all Panel I marker genes (Merge) or selected markers of distinct progenitor types in the pancreas at E12.5 and E14.5. Scale bar, 100μm. (**D**) SSAM annotated cell maps of E12.5 and E14.5 Panel I sections. Colours represent single-cell clusters as in (B). Scale bar, 100μm. (**E**) Representative HybISS images showing all Panel II marker genes (Merge) or selected markers of distinct progenitor types in the pancreas at E12.5 and E14.5. Scale bar, 100μm. (**F**) SSAM annotated cell maps of E12.5 and E14.5 Panel II sections. Colours represent single-cell clusters as in (B). Scale bar, 100μm. (**G**) Representative HybISS images of Panels I and II marker genes in E17.5 pancreas. Close-ups of selected probe genes and their spatial distribution in the tissue are shown in **i** and **ii** dotted boxes. Scale bar, 500μm.

Both panels included markers of major epithelial cell types from the pancreas: acinar, ductal, and endocrine progenitors (Fig. 1A; fig. S1). Additionally, Panel I comprised probes for endothelial, immune, neural, vascular, mesothelial, and mesenchymal cell types (Fig. 1A,C,G), while Panel II was used to explore mesenchymal cell heterogeneity (Fig. 1A,E,G). We focused on the mesenchyme, as it represents the most abundant component of the pancreatic microenvironment and is required throughout pancreas development for organ growth and cell differentiation(*10, 13–15*). Rather than being a uniform cellular population, pancreatic mesenchymal cells display heterogeneity at the transcriptomic level and, to some extent, distinct embryonic origin(*16, 17*). Consistently, we previously reported two sub-sets of mesenchymal cells (Nkx2.5- and Nkx3.2-descendant cells) showing left-right asymmetry along the embryonic pancreatic tissue and distinct functions(*18*). Thus, with the HybISS analysis, we further explored the link between transcriptomics heterogeneity, spatial location of pancreatic mesenchymal cells and associated function.

To fully assess heterogeneity within the pancreatic mesenchyme, we integrated published scRNAseq datasets(*16*) of mouse embryonic pancreatic cells with a high-coverage Smart-seq2 scRNAseq that we performed on FACS-isolated GFP^+^ cells from *Nkx2.5*-Cre;*R26*mTmG and *Nkx3.2*-Cre;*R26*mTmG transgenic (Tg) pancreatic mesenchyme and Tg(Prox1-GFP) pancreata (Fig. 1A; fig. S1A-J). Clustering identified eight mesenchymal (M) clusters with distinct gene expression profiles in line with previous studies(*16*). Two mesenchymal clusters were characterized by the expression of proliferative genes (*Mki67*, *Top2a*) (referred to as M-PI and M-PII); one cluster showed a gene signature (*Acta2*, *Tagln)* typical of vascular mural cells (M-VSM); one cluster was enriched with spleno-pancreatic mesenchymal genes, such as *Nkx2.5* and *Tlx1* (M-SpP)(*18, 19*), and one separate cluster expressed hallmark mesothelial genes (*Upk3b* and *Wt1)*(*20*). Three additional mesenchymal clusters (M-I, M-II, M-III) displayed specific marker gene sets of a less-characterized identity (fig. S1C,H) (table S2).

Pancreatic samples at different timepoints were processed for dRNA HybISS concomitantly in the same run to minimize batch effects (Fig. 1A). Out of the 96 gene probes present in the two panels we successfully detected expression of 94 of them (table S3). HybISS-based mRNA localization obtained in each dataset was used to generate spatial cell maps following two independent and complementary approaches: 1) SSAM (spot-based spatial cell–type analysis by multidimensional mRNA density estimation) segmentation-free method(*21*) and 2) segmentation-dependent method based on nuclear DAPI staining(*22–24*) (Fig. 1D,F; fig. S1K). Overall, both analyses successfully constructed spatial cell maps, which contained all cell types identified by scRNAseq, and enabled us to chart the detailed spatial relationships between these cell types and to recognize multicellular pancreatic tissue features (Fig. 1C-G; fig. S2). Notably, cell-type maps generated were comparable across all tissue sections of E12.5 and E14.5 Panel I and Panel II datasets (fig. S2).

### Mesenchymal cells display heterogeneous spatial organization across the embryonic pancreatic tissue

First, we used the HybISS-based mRNA localization to map the spatial distribution of major cell types in the pancreas and explore higher-order tissue organization in the pancreatic rudiment and to define its borders with surrounding organs, such as stomach and spleen. The pancreas forms from two distinct buds arising from the dorsal and ventral regions of the foregut endoderm, which fuse later in development to give rise to the mature organ(*6, 10*). The footprint of the two embryonic rudiments can be found in the adult tissue, wherein the dorsal pancreatic bud (DP) gives rise to part of the head, the body, and the tail of the pancreas, whereas the ventral bud (VP) to the uncinate process and the remainder of the head(*6, 10*). DP and VP progenitor cells display some differences in their transcriptional profiles both in mice and humans(*25, 26*). Here, we used HybISS spatial transcriptomics to map spatial gene expression in the DP and VP and their respective microenvironments (Fig. 2A). Within the epithelium compartment, both segmentation-dependent and SSAM analyses showed a higher number of endocrine progenitors in DP compared to VP at E12.5 and E14.5 (fig. S3A,C,E), suggesting that cellular differentiation in the developing pancreas is not uniform from a spatial point of view. In the surrounding microenvironment, we found some differences limited to the mesenchyme and the spatial distribution of transcriptionally distinct mesenchymal sub-types (fig. S3A,B). Specifically, transcripts expressed by the M-SpP (*Nkx2.5*, *Kazald1, Npnt*) were mostly restricted to the tissue surrounding the DP, whilst other mesenchymal genes in our HybISS panels (*e.g.*, *Prxx1, Pdgfrb*) were expressed around both rudiments at E12.5 (fig. S3A,B). Additionally, by HybISS and RNAScope ISH validation assay, we found differences in the expression of *Wnt5a*, a marker of the M-II sub-population (fig. S1C) (table S1), being highly enriched in the DP mesenchyme and almost undetectable in the VP mesenchyme at E12.5 and E14.5 (fig. S3C,D).

**Fig. 2.**
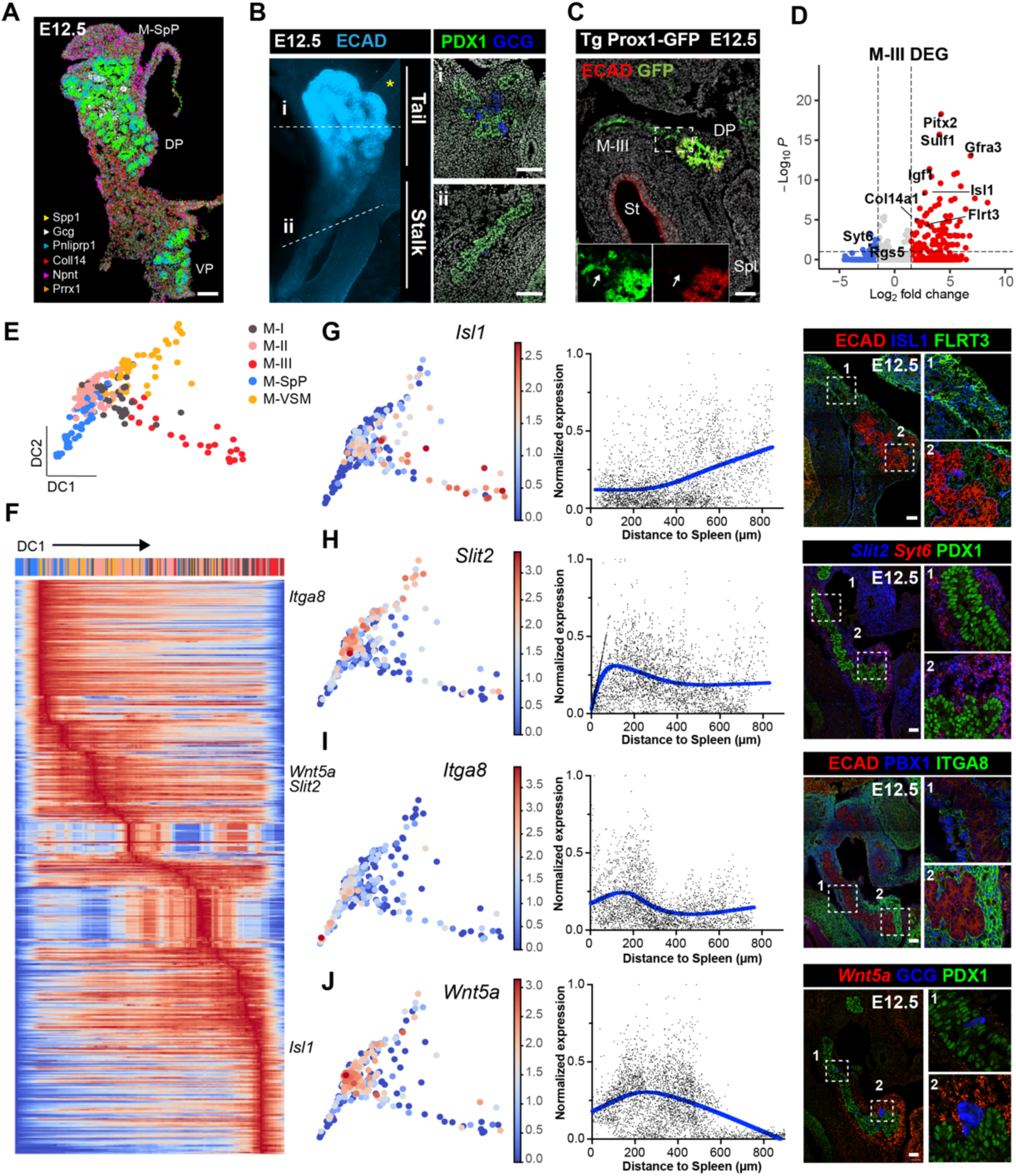
Heterogenous spatial organization of pancreatic progenitors and mesenchymal cell types across the developing pancreas. (**A**) Representative HybISS image showing selected genes from Panel I in dorsal (DP) and ventral buds (VP) at E12.5. Scale bar, 100μm. (**B**) Representative 3D-rendering of Light Sheet Fluorescent Microscopy (LSFM) image (left panel) and confocal microscopy images (right panels) of E12.5 pancreas stained with antibodies against the indicated markers. Right, confocal IF images show transverse cryosections of DP at tail (i) and stalk (ii) levels. Scale bar, 100μm. * Approximate position of the spleen. (**C**) Representative confocal image of E12.5 Tg(Prox1-GFP) DP cryosection stained with antibodies against GFP and E-Cadherin (ECAD). Arrows in the insets indicate Prox1-GFP^+^ M-III mesenchymal cells. St, stomach; Spl, Spleen. Scale bar, 100μm. (**D**) Volcano plot showing differentially expressed genes (DEG) between M-III and the other mesenchymal subpopulations. Selected top-ranked DEGs are highlighted. (**E**) Diffusion plot showing mesenchymal scRNA-seq profiles from E12.5 Smart-seq2dataset, coloured by clusters as in fig. S1A. Heatmap showing normalized and log-transformed expression levels of marker genes in the mesenchyme clusters along DC1. Genes of interest are indicated to the right. **(G-J)** Diffusion plots showing normalized and log-transformed expression levels of indicated subtype markers (left panels) in mesenchymal clusters and their validation by IF and RNAScope assays on E12.5 DP cryosections (middle and right panels). FLRT3 (G) was used as an additional marker for M-III sub-type, Syt6 (H) for M-II sub-type, and PBX1 (I) for broad DP mesenchyme (table S1). Fluorescence intensity (FI) was plotted against distance to the spleen. FI was measured with QuPATH software and values were corrected by linear normalisation within each embryo. Blue lines represent smoothing Spline curves fitting the data. Numbered dotted boxes indicate the IF magnified area shown on the right. Scale bar, 50μm.

Next, we focused on the DP, which can be subdivided into two main structural domains, based on morphological landmarks: a thin epithelium domain, connected to the duodenum (*a.k.a.* stalk) and a branched multilayered structure (*a.k.a.* tail) (Fig. 2B), which give rise to the gastric lobe and splenic lobe, respectively(*19, 27*). To assess any difference in the spatial organization of distinct cell types between the two DP domains and in their local environments, we combined HybISS spatial transcriptomics with scRNAseq and immunofluorescence (IF) analyses. Within the epithelium compartment, we found a higher number of endocrine cells in the DP tail domain when compared to the stalk, as shown by *glucagon* mRNA and protein distribution (Fig. 2B; fig. S3C,E). Interestingly, this was mirrored by the spatial arrangement of the M-SpP and M-II (*Wnt5a*-high) mesenchymal sub-populations, lying next to the tail domain, between the pancreas and the spleen (fig. S3C,D). At the opposite side of the DP, in the stalk region closer to the duodenum, mesenchymal cells mostly corresponded to the M-III cluster, positive for Prox1, and enriched for *Isl1* and *Pitx2* transcription factors, growth factors (*e.g*., *Igf1*) and ECM proteins (*e.g*., *Col14a1)* (Fig. 2C,D; fig. S3A) (table S1).

To further characterize the major axes of transcriptional variation in our data, we decided to use diffusion maps, a non-linear dimensionality reduction algorithm that orders cells along components associated with coherent gene expression patterns, preserving the underlying data structure(*28*). This method has been successfully applied to a variety of different contexts, including spatial problems(*28*). To avoid confounding factors coming from dataset integration, we focused on our deep-coverage Smart-seq2 dataset of the E12.5 mesenchyme (Fig. 2E; fig. S1B). We also removed proliferative (M-PI, M-PII) clusters to prevent cell cycle ordering being included in the dimensionality reduction. A top component of variation placed the M-SpP and M-III sub-populations at opposite sides of the inferred trajectory (pseudospace axis), with M-I, M-II and M-VSM lying in between (Fig. 2E), which closely matched the HybISS spatial data. Transcriptional variation along this diffusion component (DC1) inferred distinct patterns of gene expression across this axis, from M-SpP to M-III (referred to as splenic-gut axis) (Fig. 2F). We then used RNAscope and IF staining to spatially validate selected differentially expressed genes (DEG) enriched in distinct mesenchymal sub-types on cryosections of pancreatic tissue. Specifically, *Isl1*, whose transcript was upregulated along the first DC, displayed similar spatial distribution being abundant in M-III cells, next to the stalk region of the pancreatic epithelium, and at low abundance closer to the spleen (Fig. 2G). On the other hand, *Slit2, Itga8* and *Wnt5a* were mostly expressed in mesenchymal cells surrounding the DP tail closer to the spleen (Fig. 2H-J). These results underscored differences in the spatial organization of the mesenchyme around the pancreatic epithelium (along both dorsal-ventral and splenic-gut axes), suggesting that microenvironmental signals from the mesenchyme are responsible for the non-uniform cellular differentiation observed in the developing pancreatic epithelium.

### Topographic maps reveal dynamic endocrine and exocrine pancreatic niches

Next, we sought to use HybISS spatial transcriptomics to resolve spatial organization within the pancreatic tissue at the cellular level, including the relative positioning of different cell types and relationships among them. To this aim, we carried out two separate but complementary approaches at meso-scale (tissue-domain) or micro-scale (cell neighbourhoods) levels of organization, leveraging segmentation-free and segmentation-based frameworks, respectively (Fig. 3A).

**Fig. 3.**
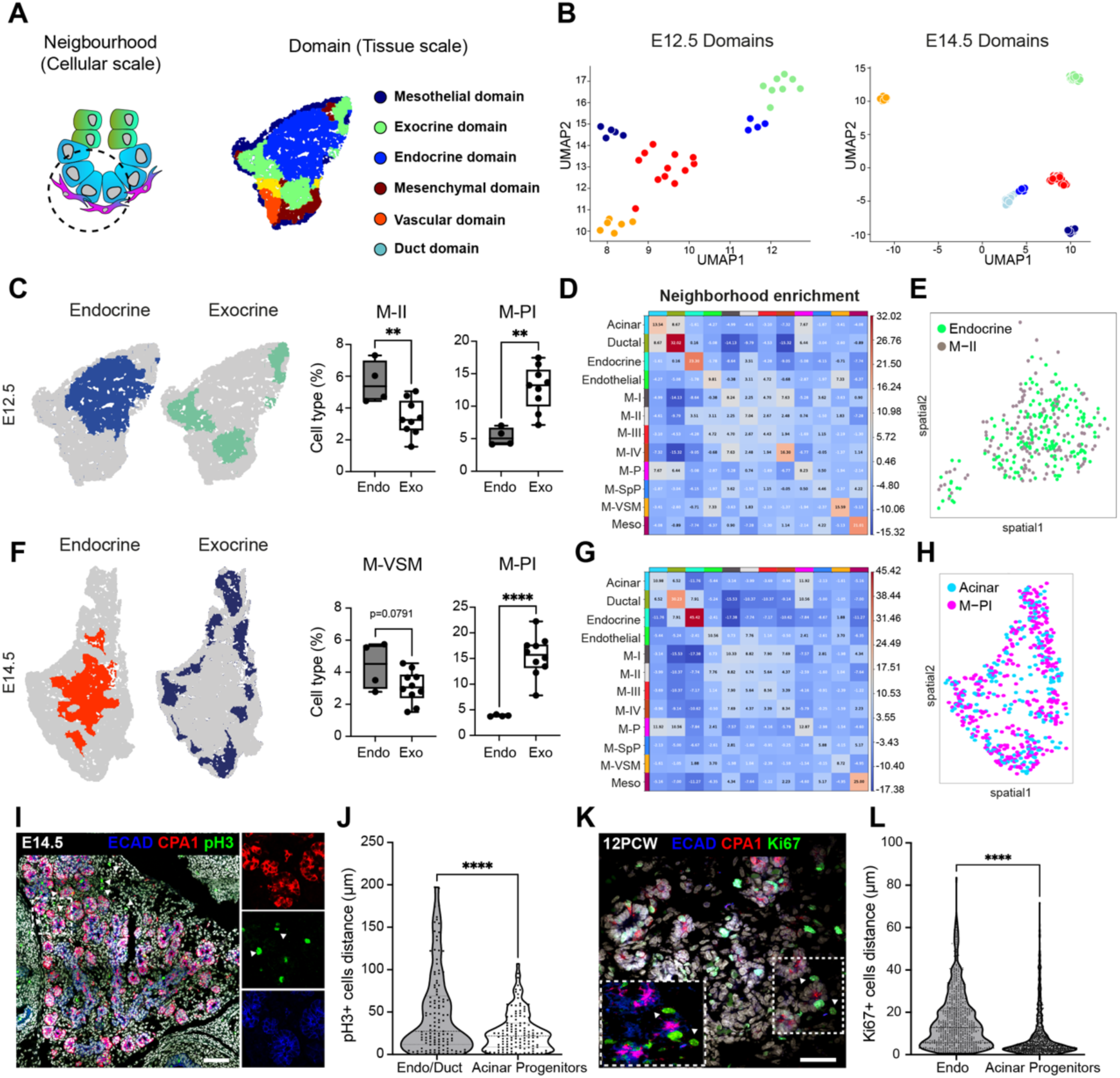
Tissue-domain analysis reveals mesenchyme niche-specific populations. (**A**) Schematics of the spatial analysis frameworks: at the cellular scale (left), spatial neighbourhoods were built by choosing the 10 closest cells around each cell and used to calculate cluster pair neighbourhood enrichment; at the tissue scale (right), tissue areas with similar local cell-type composition were clustered to identify tissue domains. (**B**) E12.5 and E14.5 tissue areas identified by SSAM on all tissue images of the Panel II dataset were grouped into specific tissue domains as shown by UMAP. Cellular composition in each tissue domain is included in table S3. (**C**) Representative endocrine and exocrine tissue domain maps of E12.5 pancreatic tissue (left) and quantification of indicated mesenchyme sub-populations in each domain (right). **p < 0.01; two-tailed unpaired t-tests. (**D**) Neighbourhood enrichment score between cell types in E12.5 pancreatic tissue images (shown as mean, n = 9 tissue sections). Positive enrichment indicates the proximity of a particular cell type to another one. (**E**) Spatial distribution of indicated cell types defined by HybISS on an E12.5 pancreatic tissue section. (**F**) Endocrine and exocrine tissue domain maps from E14.5 pancreas (left) and quantification of indicated mesenchyme sub-populations in each domain (right). ****p < 0.0001; two-tailed unpaired t-tests. (**G**) Neighbourhood enrichment score between cell types in E14.5 pancreatic tissue images (shown as mean, n = 10 tissue sections). (**H**) Spatial distribution of indicated cell types defined by HybISS on an E14.5 pancreatic tissue section. (**I**) Representative confocal image of E14.5 DP stained with antibodies against the indicated markers. Scale bar, 100μm. (**J**) Quantification of the distance between pH3^+^ proliferating mesenchymal cells to ECAD^+^/CPA1^+^ acinar progenitors or ECAD^+^/CPA1^-^ endocrine/ductal cells. n=125 cells from 3 different embryos. ****p < 0.0001; two-tailed unpaired t-test. (**K**) Representative confocal image of 12 PCW human pancreas stained with antibodies against the indicated markers. Scale bar, 50μm. (**L**) Quantification of the distance between Ki67^+^ proliferating mesenchymal cells to ECAD^+^/CPA1^+^ acinar progenitors or ECAD^+^/CPA1^-^ endocrine/ductal cells. n=748 (Endo) and n=1266 (Acinar) cells. ****p < 0.0001; two-tailed unpaired t-test.

The SSAM framework enabled us to define distinct pancreatic tissue-domains by clustering local cell-type composition from each sample in both E12.5 and E14.5 panel I and panel II (Fig. 3A-B; fig. S2) (table S4). In panel I, we identified five different tissue domains at E12.5 and E14.5 (fig. S4A-B), which were named based on the predominant cell types in each domain. Similarly, the domains in the panel II dataset were classified as exocrine, endocrine, mesothelial, mesenchymal, ductal and vascular (Fig. 3A-B and table S4). At E14.5, we observed an additional domain, primarily consisting of ductal and endocrine progenitor cells, indicating an increase in the complexity of the architecture of the embryonic pancreas (Fig. 3B). Interestingly, the cellular composition of the domains revealed a differential local enrichment of mesenchymal sub-types. For instance, at E12.5, M-PI was enriched in the exocrine domain, whereas M-II was more abundant in the endocrine domain (Fig. 3C). At E14.5, we observed the same enrichment of M-PI cells in the exocrine domain and a tendency for M-VSM to be enriched in the endocrine domain (Fig. 3F). To refine the granularity of the tissue domain analysis at micro-scale and validate its findings, we performed a neighbourhood enrichment analysis on our segmented tissue maps (fig. S2E). Consistently, M-PI was found significantly enriched in proximity to the acinar progenitors, whereas M-II was the only mesenchymal sub-type with a positive enrichment value in the endocrine neighbourhood (Fig. 3D-H). Neighbourhood enrichment analysis also validated the proximity between NCC and endocrine cells from panel I (fig. S4C-E). Finally, we validated the predicted spatial proximity between proliferative mesenchyme sub-type and acinar cells by IF staining and quantitative measurement of the distance between mesenchymal cells positive for the mitotic marker phosphorylated histone H3 (pH3) and acinar progenitor marker (CPA1) (Fig. 3I-J).

Next, to start assessing the extent of conservation of the distinct mesenchyme sub-populations and their spatial organization in humans, we first combined our integrated mouse scRNAseq dataset with a recently published human dataset(*25*) from human foetal pancreas at equivalent development stages (fig. S5A-B). The resulting dataset showed a clear resemblance between a sub-set of human mesenchyme clusters and mouse sub-populations, as well as conservation of relevant markers, ligands and ECM components in human cells (fig. S5C-D). The combined dataset was used throughout the study to assess similitudes and differences between mice and human populations. Additionally, to characterize the conservation of the acinar-M-PI spatial proximity, we performed IF analyses on human foetal pancreatic tissue at 12 post-conception weeks (PCW) using a similar set of antibodies to visualize proliferating cells in the mesenchyme (E-cadherin-negative) and CPA1-positive (+) cells (Fig. 3K). Quantitative measurement of cell-to-cell distances showed significant proximity between proliferating mesenchymal cells and acinar progenitors in human foetal pancreatic tissue, like in the mouse (Fig. 3L). Overall, our spatial analysis unveiled differential multicellular composition around endocrine and exocrine progenitor cells, highlighting mesenchymal niches with possibly distinct functional supporting role around the two main pancreatic units.

### M-II signals and secreted ECM components regulate endocrine differentiation

To identify cell-cell signalling events that might regulate pancreatic endocrine and exocrine development, we assessed the main signalling signatures across the pancreatic mesenchyme and used CellChat(*29*) to probe ligand-receptor (L:R) signalling events enriched between the identified mesenchyme sub-types and different pancreatic progenitor populations (Fig. 4A; fig. S6A-B). This highlighted a set of signalling pathways that might be implicated in pancreatic mesenchyme diversification; a sub-set of these patterns were experimentally validated (fig. S6B-F). For instance, FGF/ERK signalling was found to be mainly released by the mesothelium (source) and received by M-SpP, M-II and M-PI (targets) (fig. S6C-D). Interestingly, the transcriptome analysis of pancreatic tissue from Fgf9 knockout (KO) mice(*30*) showed gene expression modulation in mesenchymal cells along the splenic-gut axis, from M-SpP to M-III (fig. S6G-H). WNT signalling was primarily active in the epithelium (trunk progenitors), as evidenced by the quantification of LEF1 downstream effector (fig. S6 D-F). On the other hand, heterogeneous nuclear active YAP expression was detected throughout the mesenchyme (fig. S6C-D).

**Fig. 4.**
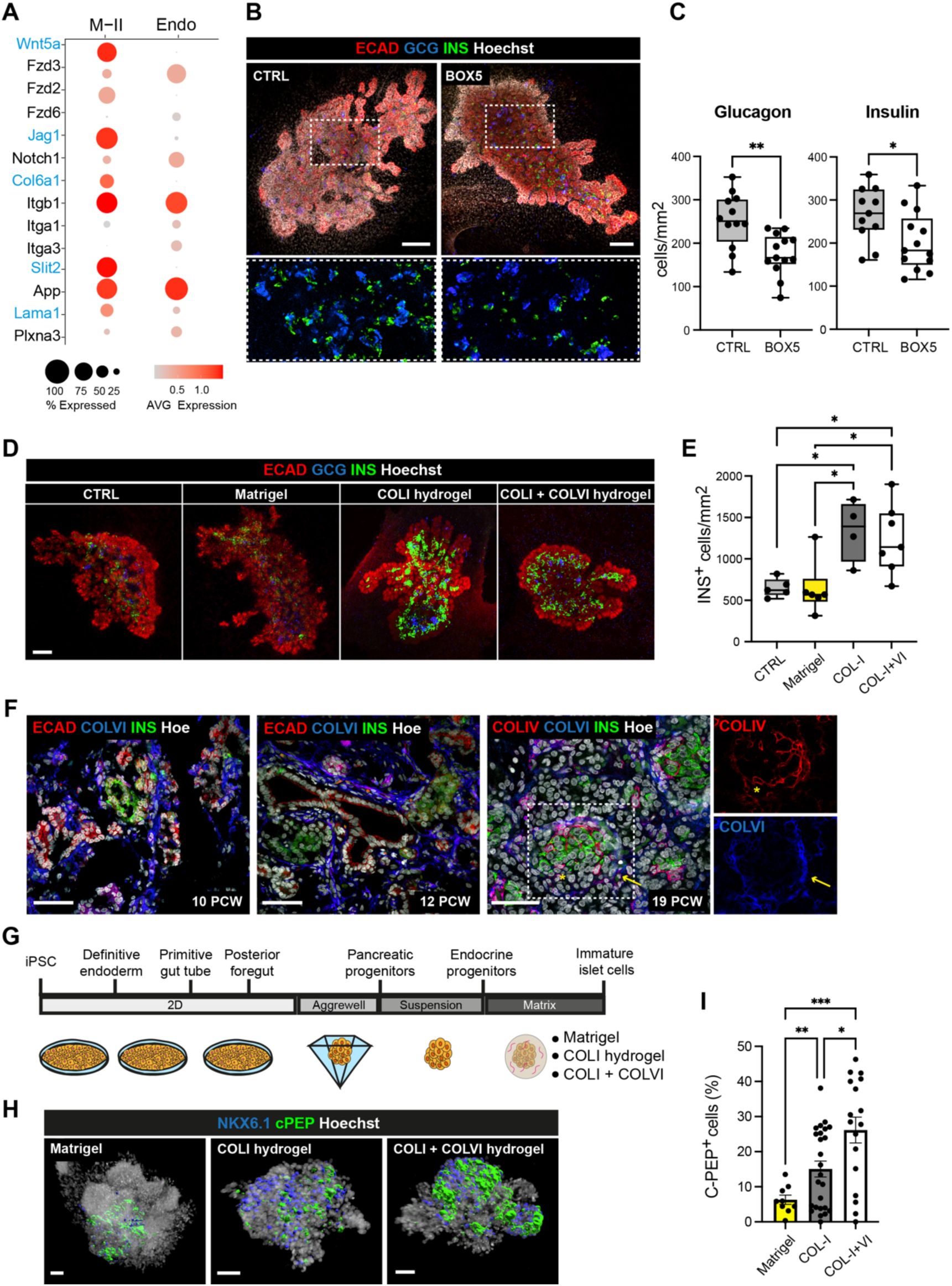
M-II signaling niche promotes endocrinogenesis. (**A**) Dot plot showing expression of ligands and cognate receptors between M-II and Endocrine cell clusters. The analysis is based on the high-coverage, Smart-seq2 dataset. Ligands are marked in blue. The colour bar indicates the linearly scaled mean of expression level. (**B**) Representative whole-mount IF images of mouse pancreatic explants treated for 2 days with BOX5 or left untreated as controls (CTRL) stained with antibodies against E-cadherin (ECAD), glucagon (GCG) and insulin (INS). Hoechst dye was used as nuclear counterstain. Bottom panels show higher magnifications of the boxed region. Scale bar, 150 μm. (**C**) Quantification of GCG- and INS-positive cells in BOX5-treated and untreated pancreatic explants. Cell counts were normalized to the average Ecad+ epithelium area (mm^2^). n=11-13 explants per condition. *p < 0.05, **p < 0.01; Two-tailed paired t-tests. (**D**) Representative whole-mount IF images of mouse pancreatic explants embedded in Matrigel, Collagen (COL) I, COLI and COLVI mix, or left unembedded (CTRL), stained for ECAD, GCG and INS. Scale bar, 150 μm. (**E**) Quantification of INS-positive cells in embedded and unembedded pancreatic explants. Cell counts were normalized to the average ECAD+ epithelium area (mm^2^). n=5-6 explants per condition. *p < 0.05; One-way Anova test. (**F**) Representative confocal IF images of human fetal pancreatic tissue between 10 to 19 PCW stained with antibodies against indicated markers. Right panels show higher magnifications of the boxed region in 19 PCW as single channels. Hoechst was used as nuclear counterstain. Scale bars, 50μm. (**G**) Schematic representation of the directed differentiation protocol of iPSCs into β-like cells adapted from(*47, 48*). iPSC-derived endocrine progenitor clusters were embedded in different matrixes at day 18 and further cultured for 4 days. (**H**) Representative whole-mount IF images of differentiated endocrine clusters at the end of differentiation stained for NKX6.1 and C-peptide (C-PEP). Hoechst was used as nuclear counterstain. C-PEP and NKX6.1 stainings were rendered as Imaris 3D surfaces. Scale bars, 50μm. (**I**) Cell quantification in 3D clusters was performed using the ‘Spots’ function in Imaris. C-PEP^+^ cell counts were normalized to the total cell number. n=25 clusters for COL-l I; n=17 clusters for COL-I + COL-VI; n=9 clusters for Matrigel. *p < 0.05, **p < 0.01, *** p < 0.001; Brown-Forsythe and Welch Anova test.

Next, given the spatial proximity identified between the endocrine and M-II cells, we focused on the crosstalk between these two populations (Fig. 4A; fig. S7A). Our predicted L:R interactions indicate strong intercellular communication between endocrine and M-II cells through the Notch signalling (Notch1, Jag1), WNT signalling (Wnt5a, Fzd3) as well as ECM molecules (Lama1, Col6a1) and integrin receptors (Fig. 4A). Some of these cell-cell communication pathways have been studied for their role in endocrinogenesis, such as the Notch pathway(*31, 32*), giving high confidence to our L:R predictions. Additionally, components of the non-canonical WNT pathway have been previously involved in regulating endocrine cell differentiation(*33, 34*), even if their spatio-temporal activities and cellular source have not been resolved. Our spatial analysis identified a localized expression of *Wnt5a* in the mesenchyme, mostly in the M-II subtype, next to the DP tail, which corresponded to the preferential site for endocrine cell differentiation (Fig. 2J; fig. S3C-D). To functionally interrogate the role of WNT5a in the pancreatic mesenchyme, we used pancreatic explants dissected from E12.5 wild-type mouse embryos, as an epithelium-mesenchyme co-culture system(*35*). Explants were cultured *ex vivo* for 48 hours in the presence of the WNT5a antagonist, BOX5, previously shown to attenuate WNT5A-mediated Ca^2+^ and PKC signalling(*36*). We found that BOX5 treatment reduced the number of insulin- and glucagon-positive cells compared to non-treated control (CTRL) samples (Fig. 4B-C). Consistently, the number of Neurogenin 3 (NGN3)-positive endocrine progenitors was also decreased in BOX5-treated pancreatic explants (fig.S7B-C). Together, these results are in support of a role for WNT5A in endocrine cell differentiation.

Next, we sought to investigate the ECM molecules that compose the M-II niche and their potential impact on pancreatic endocrine development. We primarily focused on Collagen VI (COLVI), because despite being among the most abundant ECM molecules in the adult pancreatic islets(*37*), its function in the context of pancreatic development is unknown. We found that COLVI increases, both at the transcript (*Col6a1*, *Col6a2* and *Col6a3*) and protein levels, during development (fig. S7D-E), being densely distributed around the endocrine progenitors and insulin-positive cells in the mouse and human foetal pancreas, respectively (Fig. 4F; fig. S7E). To investigate the role of the ECM composition on pancreatic development, we embedded mouse pancreatic explants in hydrogels containing different ECM components and assessed growth, morphology and differentiation. Both COLI and COLI/COLVI (*i.e.,* interstitial-like ECM) led to an increase in the number of insulin-positive cells compared to Matrigel (*i.e.*, basal membrane-like ECM, including mostly COLIV and laminins) and non-embedded controls (Fig. 4D-E), whilst the number of glucagon-positive cells was unchanged (fig. S7F). Murine pancreatic explants also showed differences in their overall morphology and size according to the hydrogel composition. Explants embedded in Matrigel underwent extensive branching, whilst the ones in Collagen hydrogel showed a rounder morphology (Fig. 4D). Next, to study the functional conservation of the ECM effect in human endocrine differentiation, we used a modified human pluripotent stem cell culture system for modelling human pancreatic development (Fig. 4G). Specifically, iPSC cells were first differentiated to the pancreatic progenitor stage in 2D, then embedded at the endocrine progenitor stage in the same hydrogel composition used for the mouse explants and cultured for 5 days, until collection (Fig. 4G). In human cells, COLVI had a specific beneficial effect, promoting an increase in the number of insulin-positive cells as compared to the COLI alone and Matrigel hydrogels (Fig. 4H-I). Taken together, these findings unveiled an endocrine niche, enriched for M-II mesenchyme, with conserved composition, spatial organization and function in human.

### Embryonic Nkx2-5^+^ M-SpP progenitors give rise to distinct populations of adult pancreatic fibroblasts

Mesenchymal progenitors give rise to mature adult tissue-resident fibroblasts, which support organ homeostasis and participate in fibrosis and diseases, like cancer(*38*). Despite adult fibroblasts playing a major role in pancreatic fibrosis and cancer progression and reactivating developmental programs at the injury site to some extent(*39–41*), their ontogeny and lineage relationships in the embryonic pancreas are still elusive. To answer these questions, we extended our analysis to adult tissue homeostasis. First, we integrated scRNAseq datasets of healthy mouse pancreatic tissue from three different studies(*3, 42, 43*) (Fig. 5A-B). After batch correction and clustering we identified a mesothelial cluster (Krt18^+^), a vascular cluster (Acta2^+^) and two main adult fibroblast clusters (Fig. 5A-B). Adult fibroblast-I (AF-I) cells expressed genes that have been previously linked to reticular fibroblasts (*i.e*., *Klf4*, *Mfap5*, *Cd34)*(*44*), while AF-II cells were characterized by a high level of expression of Collagen and other ECM genes, including *Col6* genes, and high levels of *Smoc2* (Fig. 5C,F). Next, we quantified the distribution of both AF-I and AF-II in different niches of the adult pancreas. Both cell types were similarly represented in the acinar and ductal niches but showed differences with respect to the islets, being both at the islet periphery and only AF-I cells infiltrating them (Fig. 5D-E). Finally, to understand the developmental origin and build a cellular taxonomy of the mesenchymal lineage from embryonic pancreas to adult homeostasis, we used *in silico* pseudotime and *in vivo* genetic lineage tracing experiments. First, we integrated embryonic and adult mesenchymal sc-datasets (Fig. 5G). Then, we used Slingshot to obtain lineage trajectories and projected them into the UMAP representation (Fig. 5G). Notably, the M-SpP (*Nkx2.5^+^*) mesenchyme was ordered at the origin of the trajectories with different trajectories for the vascular cells (A-VSM) and the two AFs. The lineage bifurcation of adult fibroblasts started during embryonic development with M-III giving rise preferentially to AF-I cells and the other embryonic clusters to AF-II (Fig. 5G). Next, we used the *Nkx2.5*-Cre mouse Tg model with the *R26*-mTmG reporter line to track embryonic mesenchymal cells into adult tissue. In line with the pseudotime model, our analysis revealed the presence of KLF4^+^ GFP^+^ (AF-I), SMOC2^+^ GFP^+^ (AF-II) and SMA^+^ GFP^+^ (AM-VSM) cells in the adult pancreas (Fig. 5H-I), confirming the contribution of embryonic fibroblasts to adult pancreatic stroma.

**Fig. 5.**
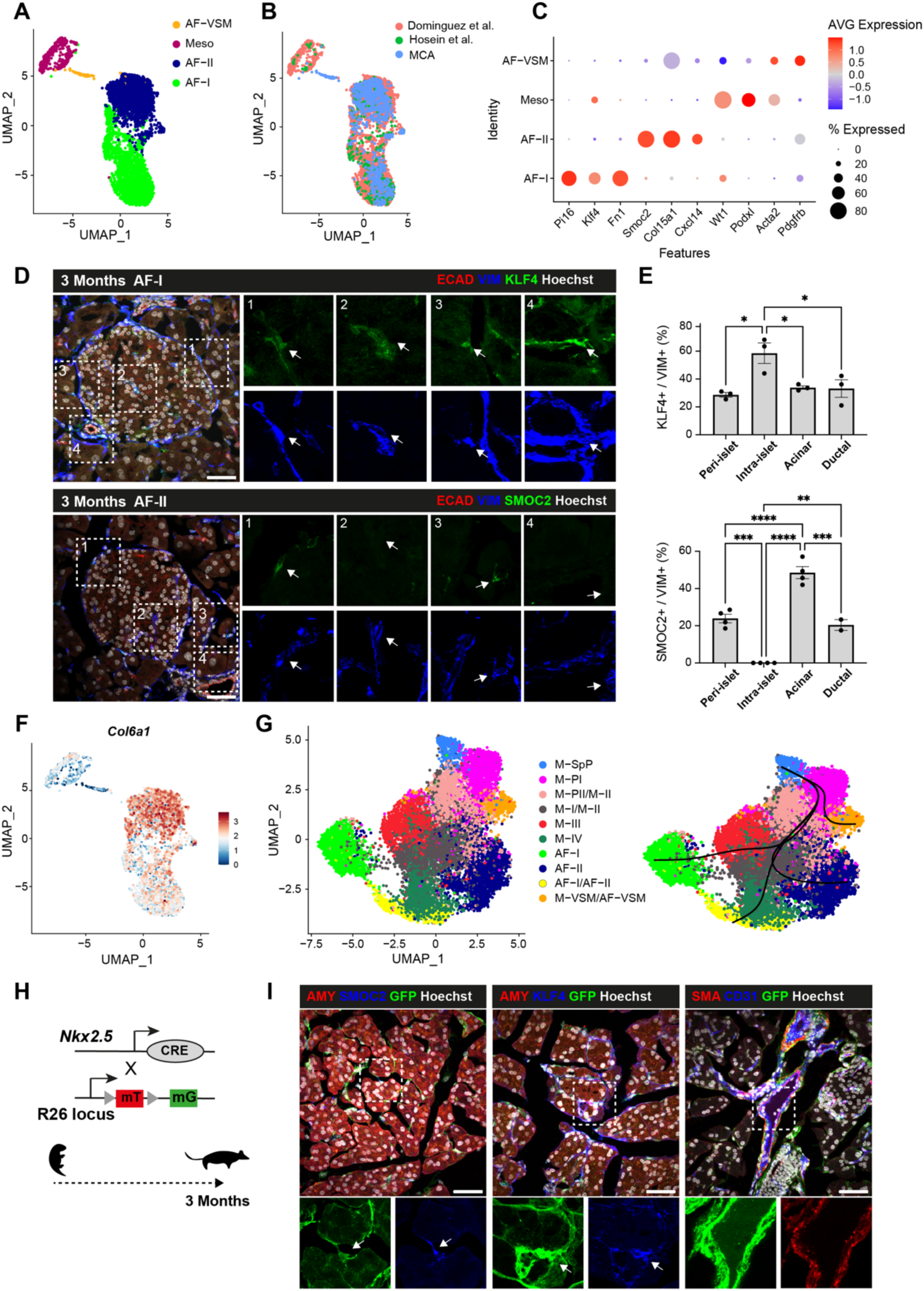
Embryonic Nkx2.5^+^ M-SpP lineage contributes to adult pancreatic fibroblasts. (**A**, **B**) UMAP plot visualization of fibroblast cells in the adult pancreas from 3 integrated sc-RNAseq datasets(*3, 42, 43*). Clusters are distinguished by different colours, based on cell type identity using known marker genes (A) or dataset (B). (**C**) Dot plot showing the expression of selected marker genes for each cell cluster from (A). Colour bar indicates the linearly scaled mean of expression level (see table S4). (**D**) Representative confocal images of adult pancreas cryosections stained with antibodies against the pan-mesenchymal marker Vimentin (VIM) and adult fibroblasts (AF)-I-specific KLF4 and AF-II-specific SMOC2 markers. Right panels show higher magnifications of the boxed regions as single channels. Arrows indicate AF-I (KLF4^+^) and AF-II (SMOC2^+^) cells in contact with specific epithelial structures (1: Peri-Islet; 2: Intra-Islet; 3: Acinar; 4: Ductal). Hoechst was used as nuclear counterstain. Scale bars, 50μm. (**E**) Quantification of the percentage of SMOC2+ and KLF4^+^ cells in each niche over the total VIM+ cells. n=3-4 pancreas per staining. Error bars represent ± SEM. *p < 0.05, **p < 0.01, **p < 0.01, *** p < 0.001, **** p < 0.0001; One-way Anova test. (**F**) Feature plot showing *Col6a1* expression in adult pancreatic fibroblast cells. Colour bar represents mean log-transformed expression. Clusters are labeled based on the annotation in Fig. 5A. (**G**) Left, UMAP plot showing clustering of sc-RNAseq profiles of mesenchymal cells from embryonic and adult pancreas integrated dataset. Clusters are labeled based on the annotation of adult (as in Fig. 5A) and embryonic mesenchyme datasets (as in Fig. 1B). Right, slingshot-inferred lineages on the embryonic-adult mesenchyme clusters, showing putative hierarchy of the pancreatic mesenchymal lineage. (**H**) Schematic representation of the lineage tracing strategy using *Nkx2.5*-Cre;*R26*mTmG mouse model. Recombination occurs in embryonic tissue, whereby Nkx2.5-expressing-M-SpP mesenchymal cells acquire GFP (mG) label and are then tracked in adult pancreas. (**I**) Representative confocal images of IF staining on cryosections of *Nkx2.5*-Cre;*R26*mTmG adult pancreas for the indicated markers. Bottom panels show higher magnifications of the boxed regions as single channels. Arrows indicate *Nkx2.5*-Cre-descendant cells contributing to distinct adult fibroblast populations. Hoechst was used as nuclear counterstain. Scale bars, 50μm.

## Discussion

Our study provides a high-resolution spatiotemporal profiling of the developing pancreas, resolving the spatial organization of pancreatic cells and their surrounding microenvironment at multiscale level. We identified potential mediating pathways for cell proximity effects at the organ scale, as well as the multicellular mechanisms that define spatial niches within the two main functional compartments of the pancreas: the endocrine and exocrine domains. Notably, these niches displayed differences in the mesenchymal sub-type composition; the M-II was enriched next to endocrine progenitors, while the proliferative M-PI next to acinar progenitors. These findings extend previous work that has underscored transcriptomic heterogeneity within the pancreatic mesenchyme(*16, 17, 25*), demonstrating that mesenchymal cells comprise several distinct, functional sub-types, distinguishable based on their gene expression profiles, origin and location. Although we identified signaling pathways establishing spatial gradients in the mesenchyme, more functional studies will be needed to provide a better understanding of their role in mesenchymal cell fate diversification.

Here, we focused on disentangling the proximity effect of heterogeneous mesenchymal cell populations in different regions of the tissue. For instance, we discovered that the M-II sub-type is a source of a COLVI-rich ECM, and we demonstrated its pro-endocrine activity, a feature conserved in both mouse and human. Interestingly, COLVI has been previously identified as a major component of the islet-exocrine interface in the adult pancreas(*37, 45*); here, we shed light on its role in the embryonic pancreas and, specifically, in endocrine differentiation. Typically, COLVI functions *via* direct engagement of cell surface receptors, such as integrins, but it also has the means to sequester cell-signalling ligands, such as PDGF, as well as to regulate the mechanical properties of the cell microenvironment(*46*). Additional studies will be needed to provide a better understanding of the mode of action of COLVI and its biological effects in the pancreas. Ultimately, broader investigation into cell spatial organization, cell-cell interaction effects and their mediators will be critical and may lead to new therapeutic strategies, improving the engineering of complex multicellular tissues for human pancreatic beta-cells. Finally, our study defined the contribution of embryonic mesenchyme to the adult pancreatic stroma underscoring the possibility that changes in the mesenchyme composition or niche spatial organization during fetal life might play a role in pancreatic disease. Expanding systematic spatial investigations will be critical to assess whether such deleterious alterations of the endocrine niche in embryonic life could contribute to diabetes pathogenesis.

## Acknowledgement

We thank all the members of the Spagnoli laboratory for their useful comments and suggestions on the study. We thank Heiko Lickert (Helmholtz, Munich) for the hiPSC line HMGUi001-A2. We thank Christian Beisel of the Genomics Facility of D-BSSE for NGS RNA sequencing. We thank the Human Developmental Biology Resource (http://hdbr.org) for the help in collecting the pancreatic foetal tissue. We are grateful to the BRC Flow Cytometry platform at King’s College London.

## Author contributions

F.M.S. and A.T.C. conceived the study, designed the experiments and wrote the manuscript with input from remaining authors. A.T.C. performed all the experiments. J.F.D. and A.T.C. designed and performed the HybISS experiments together with M.R. and D.C. G.G. performed and analysed all experiments in human foetal tissue. D.W. helped with mouse work and collection of cells for scRNAseq. A.S. performed the light-sheet microscopy image acquisition. A.T.C., A.V., G.E.H. conducted sc-RNASeq data and spatial HybISS analyses.

## Data and materials availability

Smart-seq2 sc-RNAseq and HybISS spatial transcriptomics data generated in this study have been deposited in the Gene Expression Omnibus (GEO) under accession number GSE283594 and GSE290835, respectively. Public sc-RNAseq data were collected from GEO (GSE101099, GSE125588, GSE176063) and from ArrayExpress database (E-MTAB-8483). All data are available in the main text or the supplementary materials. Correspondence and request for materials should be addressed to FMS.

## Supplementary Materials

### Materials and Methods

#### Mouse Strains

All procedures relating to animal care and treatment conformed to the Institutional Animal Care and Research Advisory Committee and local authorities (PPL PP6073640, Home Office, UK). All mouse embryos were used without sex identification (mixed sexes). The following mouse strains were used in this study: Nkx2-5^tm2(cre)Rph^(*49*), Nkx3-2^tm1(cre)Wez^(*50*), Tg(Prox1-EGFP)(*26*), B6.129(Cg)-Gt(ROSA)^26Sortm4(ACTBtdTomato,-EGFP)Luo/J^(*51*). All mice were bred on a C57BL/6J genetic background. Mice were housed in a specific pathogen-free facility in individually ventilated cages. Room temperature was maintained at 22±1°C with 30–70% humidity and lighting followed a 12-h light/dark cycle. Food and water were provided ad libitum and none of the mice had been involved in previous procedures before the study. For timed mating, male and female mice were placed into a breeding cage overnight (ON) and plug check was performed daily. The presence of a vaginal plug in the morning was noted as E0.5.

#### Pancreatic Explants Culture

Dorsal pancreatic buds were microdissected from mouse embryos at E12.5 and cultured on glass-bottom dishes (Matek) pre-coated with 50 mg/ml sterile bovine Fibronectin (Invitrogen) in Basal Medium Eagle (BME) (Sigma) supplemented with 10% FBS (Invitrogen), 1% Glutamine, 1% Penicillin-Streptomycin (PS) (Invitrogen)(*35*). The day of plating is referred to as day 0. Explants were cultured in a tissue incubator (37°C, 5%CO2) and culture medium was changed daily. For WNT5a blocking experiment, BME culture medium was supplemented after 24h with BOX5 (50 μg/mL) (Sigma). For matrix experiments, at day 2 of culture, the medium was removed, and the explants were overlaid with the matrix solutions, which were allowed to polymerize for 30 minutes at 37°C. BME media was then added to the plate and the explants were cultured for 3 additional days. Collagen hydrogel solutions were prepared by mixing Collagen I (Serva) with or without Collagen VI (Abcam), BME media, 10X PBS and neutralized using NaOH. Matrigel mixture was prepared by mixing BME media and Matrigel (Corning) in 1:1 proportion. At the end of the treatment, the explants were briefly washed with PBS, fixed for 20 minutes at 4°C in 4% paraformaldehyde (PFA) and processed for whole-mount immunofluorescence (IF) as previously described(*43*).

#### Immunohistochemistry and RNAscope

Mouse embryos and pancreata were fixed in 4% PFA at 4°C from 2 hours to ON. Human embryonic and foetal tissue was provided by the joint MRC/Wellcome Trust (grant# MR/X008304/1 and 226202/Z/22/Z) Human Developmental Biology Resource (http://hdbr.org) with appropriate maternal written consent and approval from the Newcastle and North Tyneside NHS Health Authority Joint Ethics Committee (23/NE/0135) and London Fulham Research Ethics Committee (23/LO/0312). The HDBR is regulated by the UK Human Tissue Authority (HTA; www.hta.gov.uk) and operates in accordance with the relevant HTA Codes of Practice. Human pancreatic tissue samples (gender not established) were fixed overnight in 4% PFA and then processed for cryosectioning, immunostaining and imaging at King’s College London. All work was undertaken in approval of the HDBR Steering Committee to the Spagnoli lab. at King’s College London, UK (License #200523).

For cryosectioning, samples were equilibrated in 20% sucrose solution, embedded in OCT compound (Tissue-Tek, Sakura) and sectioned at 10μm thickness. For immunostaining, sections were incubated in TSA (Perkin Elmer) blocking buffer for 1h at room temperature (RT). If necessary, antigen retrieval was performed by boiling slides for 20 min in citrate buffer (Dako). Sections were incubated in primary antibody solution (3% horse serum and 3% BSA in PBS) at the appropriate dilution (see table S7) ON at 4°C. Hoechst 33342 nuclear counterstaining was used at a concentration of 250 ng/mL. For RNAscope *in situ* hybridisation, cryosections were processed with the RNAscope Multiplex Fluorescent Reagent Kit v2 (Advanced Cell Diagnostics) according to the manufacturer’s instructions. Protease Plus (diluted 1:5) was applied to permeabilise samples for 10 minutes. Probes (see table S7) were used in combination with Opal 650 and Opal 570 fluorophores. Images were acquired with Zeiss AxioObserver, Zeiss Discovery and Zeiss LSM 700 laser scanning confocal microscopes.

#### Tissue clarification and light-sheet microscopy

Whole-mount IF staining on pancreatic tissue was performed as previously described(*52*). After whole-mount staining, tissue clarification was performed using CUBIC1 [25% wt/vol urea, 25% wt/vol N,N,N′,N′-tetrakis(2-hydroxypropyl) ethylenediamine, 15% wt/vol Triton X-100, in dH2O] and CUBIC2 (50% wt/vol sucrose, 25% wt/ vol urea, 10% wt/vol 2,20,20′-nitrilotriethanol, 0.1% vol/vol Triton X-100, in dH2O) solutions. After clarification, samples were glued to a 1 ml syringe supportive holder using all-purpose super glue. The syringe was then inserted into the syringe holder provided with the Zeiss Z1 light-sheet microscope. Samples were then imaged with the Zeiss Z1 light sheet microscope using 20X acquisition and 10X illumination lenses.

#### Library preparation and RNA sequencing

Pancreatic tissue from E12.5 Tg(Prox1-EGFP), Nkx2.5-Cre;R26mTmG and Nkx3.2-Cre;R26mTmG embryos were manually dissected in DEPC-PBS, individually digested in Collagenase for 10 minutes at 37 °C and processed for FACS sorting, as previously done in(*53*). Tissue digestion was blocked by adding DMEM medium with 10% FBS, cells were spun for 3 minutes at 300g at 4 °C, resuspended in DEPC-PBS and filtered through FACS tubes with Cell Strainer Cap (BD 352235) for immediate FACS sorting (BD FACS Aria II or III). Single cells were sorted into 96-well plates in lysis buffer (0.2% Triton X-100, 2 U/μl Rnasin in nuclease-free water). Library preparation and transcriptome sequencing were performed by the Genomic sequencing facility at D-BSSE (Basel), according to the Smart-seq2 protocol(*53*). Sequences were obtained from E12.5 GFP positive cells from Nkx2.5-Cre;R26mTmG (192 cells), Nkx3.2-Cre;R26mTmG (192 cells), and Tg(Prox1-EGFP) (193 cells). cDNA profiles were checked on the Fragment Analyzer (AATI) and their concentration determined using Quant-iT PicoGreen dsDNA Assay Kit. Libraries were pooled and sequenced SR75 on an Illumina NextSeq 500 system (75 cycles High Output v2.5 kit).

#### Sc-RNAseq processing

Reads were uniquely aligned to the mouse reference genome (GRCm38) using Kallisto pseudo-mode (v.0.46.1). The gene expression matrices were generated by converting transcript level counts to gene-level counts using Tximport (v.1.18.0).

Downstream analyses were completed with Seurat (version 4.4.0)(*54*). Quality control was performed to remove genes expressed by less than three cells, exclude cells expressing fewer than 1000 genes, less than 100000 gene counts, or a percentage of counts derived from the mitochondrial genome higher than 30%. Data was Log Normalized and cell cycle differences were regressed using the ScaleData function. We used Principal component analysis (PCA) for dimension reduction and unsupervised clustering of the data with the FindNeighbors() and FindClusters() functions. Meaningful PCAs were selected using the Jackstraw function and used for the FindNeighbors() function to construct a shared nearest-neighbour graph of all the data. Then, the cells were clustered using the function FindClusters() with a shared nearest-neighbour modularity optimization-based clustering algorithm with a resolution of 1.9. To visualize the data, we used uniform manifold approximation and projection (UMAP). Clusters were finally interrogated and manually curated based on their differentially expressed genes. E12.5, E14.5 and E17.5 datasets (GSE101099)(*16*) were obtained from the Gene Expression Omnibus (GEO) database. Quality control was performed using values specified by the authors. To integrate it with our Smart-seq2 dataset, each separate batch was pre-processed independently using the SCTransform() function(*55*) with 3.000 features and regression of differences in cell-cycle state among cells. Integration was then performed using the Seurat integration workflow. Downstream processing was performed as previously specified. The first 20 PCA were used for the FindNeighbors() functions and a resolution of 1 was used to find clusters.

Adult scRNA-seq datasets were obtained from published datasets GEO (GSE125588, GSE176063)(*3, 43*) and ArrayExpress database (E-MTAB-8483)(*42*). Quality control was performed using values specified by the authors. For integration, each separate batch was pre-processed independently using the SCTransform() function(*55*) with 3.000 features and regression of differences in cell-cycle state among cells. Integration was then performed using Seurat integration as previously specified. The first 30 PCAs were used for the FindNeighbors() functions and a resolution of 0.7 was used to find clusters. For adult and embryonic dataset integration, adult and embryonic mesenchymal cells were integrated using the Harmony package(*56*). The first 50 PCAs were used for the FindNeighbors() functions and a resolution of 0.5 was used to find clusters.

Human scRNA-seq data was obtained from the OMIX database (OMIX001616)(*25*). Quality control was performed using values specified by the authors. After subsetting mesenchymal cells from each batch, integration was performed using Seurat workflow as previously specified. For human-mouse integration, mouse orthologues were first mapped to human genes using the Python package Mousipy (v.0.1.6). Next, human and mouse datasets were integrated using MultiMAP(*57*) using separately pre-calculated principal with 0.7 and 0.3 strength respectively. Differentially Expressed Genes (DEG) were found using the Seurat’s FindAllMarkers() Function.

#### Ligand-Receptor interaction analysis

CellChat (v.1.6.1)(*29*) was used to investigate cellular interactions based on our Smart-seq2 dataset. First, we used the CellChatDB ligand-receptor interaction database and then the CellChat pipeline to compute the interaction number and strength between the different cell types as well as the incoming, the outgoing signalling patterns and the specific ligand-receptor interactions using default parameters.

#### Diffusion analysis

To perform diffusion analysis on our Smart-seq2 dataset, we used Scanpy (v.1.9.0)(*58*). First, we excluded epithelial, mesothelial and proliferating mesenchymal cells. Then, we generated a normalized, log-transformed cell-by-gene matrix of 2000 variable genes which we used to generate diffusion maps using the Scanpy tl.diffmap() function. Meaningful diffusion components were used to generate a diffusion plot, which we subsequently used to visualize cell identities, gene expression ang gene module expression. To investigate gene expression trends associated with underlying transcriptional heterogeneity, we used the pyGAM package (v. 0.8.0). Specifically, we used the 1000 most variable genes and fit their gene expression along DC1. Significantly associated genes were defined as those having a p-value <0.01. Gene module analysis was performed with the GSEApy package (v.1.1.4) to retrieve gene signatures from Biocarta (namely, ‘Signaling by Hippo’, ‘ERK MAPK Targets’ and ‘Signaling by WNT’). The Scanpy tl. tl.score_genes() function was then applied to generate signature scores. For the FGF9 KO analysis, we retrieved upregulated and downregulated gene signatures from(*30*). The Scanpy tl. tl.score_genes() function was then applied to generate signature scores that were subsequently plotted in the diffusion plot.

#### Hybridization-based *in situ* sequencing (HybISS) for spatially resolved transcriptomics

E12.5, E14.5 and E17.5 embryonic pancreata were dissected and fixed in 4% PFA ON. Samples were equilibrated in 20% sucrose solution and embedded in OCT compound (Sakura). 10 μm-thick cryosections were stored at −80°C. On the starting day of the experiment, sections were incubated at 60°C for 1 hour before post-fixation with 4% PFA for 5 minutes. After washes, sections were serially dehydrated in 50%, 70% and 100% Ethanol, subjected to antigen retrieval in Citrate buffer and then hybridized with the gene panels at 37°C ON. Ligation and following steps were performed using the High Sensitivity Library Preparation Kit from Cartana AB (10x Genomics) as described(*12*). Incubations were performed in SecureSealTM chambers (Grace Biolabs, Bend, OR, USA). SlowFade Antifade Mountant (Thermo Fisher Scientific, Waltham, MA, USA) was used for optimal handling and imaging of tissue sections. After quality control samples were processed by Cartana AB (10x Genomics), for *in situ* barcode sequencing, imaging, and data processing. This generated an output consisting of DAPI images of tissue sections and CSV files, containing gene identity and position of identified RNA spots. To generate gene expression tissue plots we used MATLAB. For downstream analysis we manually generated a mask around the pancreas to filter out cells and reads from other tissues. E12.5 and E14.5 datasets were analysed by segmentation-free and segmentation-based computational frameworks, while E17.5 dataset was used only for qualitative analysis due to the limited number of sections analysed and low detection rate.

#### Cell segmentation-free approach (SSAM)

Segmentation-free analysis of the HybISS data was performed following the SSAM pipeline (v.1.0.2)(*21*). Gene reads coordinates were first transformed from pixels into micrometres (0.32 µm per pixel) and then used to generate mRNA density maps through a Kernel Density Estimation (KDE) (Gaussian kernel and a bandwidth of 2.5). Gene density maps were then combined to identify local maxima using a total gene expression threshold of 0, a per gene expression threshold of 0, and a search size of 3. Local maxima were then used to calculate the variance stabilization parameters with the SCTransform package (v0.3.4)(*55*) which were subsequently used for gene expression normalization. For cell type identification, we used the integrated dataset containing E12.5, E14.5 and E17.5 cells as the reference dataset. The raw gene counts were normalized using SCTransform and the average gene expression per cell type was calculated. Local maxima vectors were mapped to the cell type clusters with the most similar gene expression in the scRNAseq reference dataset, using a correlation minimum threshold of 0 and a threshold of vector normalisation of 0.025. The obtained tissue maps of each section were then analysed independently to build tissue domains. The combined list of tissue domains from all samples was then clustered and visualized using k-means clustering, PCA and UMAP. Tissue domains were finally manually annotated based on the most abundant cell types in each domain.

#### Cell segmentation-based approach

For the segmentation-based approach, we first used Cellpose 2.0(*23*) to segment cells based on DAPI staining. Next, we assigned reads to each cell using Probabilistic cell typing for *in situ* sequencing (PciSeq)(*24*). This generated a Cell x Gene matrix, which was then used as input for Tangram(*22*) to annotate cell type based on the integrated sc-RNAseq dataset. The resulting tissue maps were then combined and used to build neighbourhood enrichment plots using Squidpy(*59*). Squidpy (v.1.1.4) was used to compute a spatial graph with the gr.spatial_neighbors() function and a window of 10 neighbours. We then used this graph to compute neighbourhood enrichment scores.

#### Cell lines and Cell culture

Human iPS cell line HMGUi001-A2 (sex: female)(*60*) was kindly provided by Heiko Lickert (Helmholtz, Munich) and authenticated by karyotyping. Human iPSCs were maintained on Geltrex-coated (Invitrogen) plates in E8 media. The medium was changed daily and cells were passaged every 3-4 days as cell clumps or single cells using 0.5 mM EDTA (Invitrogen) or Accutase (Invitrogen), respectively. The medium was supplemented with Rho-associated protein kinase (ROCK) inhibitor Y-27632 (Sigma) (10 μM) when iPSCs were thawed or passaged as single cells.

#### Differentiation of Pluripotent iPSCs into Pancreatic β-like Cells

Human iPSC differentiation was carried out following a protocol adapted from published ones(*47, 48*) (table S8). Briefly, iPSCs were dissociated using Accutase and seeded at a density of 2.8×10^6^ cells per well in 6-well plates (Corning) coated with Geltrex (Invitrogen) in E8 medium supplemented with ROCK inhibitor (10 μM). Cells were differentiated into pancreatic progenitors as previously described(*47, 48*). At day 11 of the protocol, pancreatic progenitors were dissociated using Accutase and seeded to microwells (24-well AggreWell 400 microwell plates, Stem Cell Technologies) at a density of 1000 cells per microwell. Cells were maintained in stage 4 media in AggreWells for 3 days. On the first day of Stage 5, endocrine progenitors were transferred from AggreWells to ultralow attachment plates (CellStar) and placed on an orbital shaker for suspension culture at 100 r.p.m. At day 4 of Stage 5, 3D cell clusters were collected and resuspended in Matrigel or collagen hydrogel mixtures. The matrix was allowed to polymerize for 30 minutes at 37 degrees. Subsequently, the media was added to the plate and clusters were cultured for additional 5 days in Stage 6 media. Collagen hydrogels were prepared as described for pancreatic explants.

#### Quantifications, statistics and reproducibility

Cell numbers and immunostaining intensities were quantified on sections and explants using QuPATH(*61*). For cell quantification in embryonic samples, the entire pancreas was serially sectioned (10μm thick). For cell counting, positive cells (matching a Hoechst^+^ nucleus) were manually counted or using the QuPATH positive cell detection function. The relative area occupied by cells per pancreas was obtained by normalizing the counted immunopositive cells to the total E-cadherin^+^ or PDX1^+^ epithelium area.

For cell quantification in whole-mount pancreatic explants, immunopositive cells were counted in three focal planes of a Z-stack (top, centre and bottom) and normalized to the average epithelial area (X.Y) multiplied by the number of focal planes analysed in the explant (Z).

Analysis of mesenchymal cell positioning along the gut-splenic axis was performed by measuring IF intensities in non-epithelial cells (E-cadherin^-^ or PDX1^-^) and their distance to the spleen using QuPATH. IF intensities were corrected by linear normalization within each embryo to achieve a uniform dynamic range and improve comparability between embryos. To show the curves fitting the data we used GraphPad function Spline curves with 5 knots. Quantification of *Wnt5a* expression and LEF1 fluorescent intensity in embryonic tissue sections was performed using QuPATH. In adult samples, quantifications were performed across five to seven independent fields of view in each of three different sections per adult pancreas. For iPSC cell quantification, we used the Imaris Spot detection function. Experiments were repeated a minimum of three times; one representative field of view is shown for each staining. All results are expressed as mean ± standard deviation (s.d.) or mean ± standard errors (s.e.m), as indicated. Sample sizes of at least n=3 were used for statistical analyses except where indicated.

Data representation and statistical analysis were performed using GraphPad Prism (version 9) and Excel (Microsoft). Statistical significance was determined as indicated in figure legends using unpaired two-tailed Student’s t-test, Mann–Whitney test, or Anova test for more than two groups. Statistical analysis was conducted on all collected samples and data. No statistical method was used to predetermine the sample size. No data were excluded from the analyses. The experiments were not randomized. Investigators were not blinded to allocation during experiments and outcome assessment.

## Data availability

Smart-seq2 sc-RNAseq and HybISS spatial transcriptomic data generated in this study has been deposited in the Gene Expression Omnibus (GEO) under accession number GSE283594 and GSE290835, respectively. Public sc-RNAseq data were collected from GEO (GSE101099, GSE125588, GSE176063) and from ArrayExpress database (E-MTAB-8483). Source data are provided with this paper.

---

**Fig. S1.**
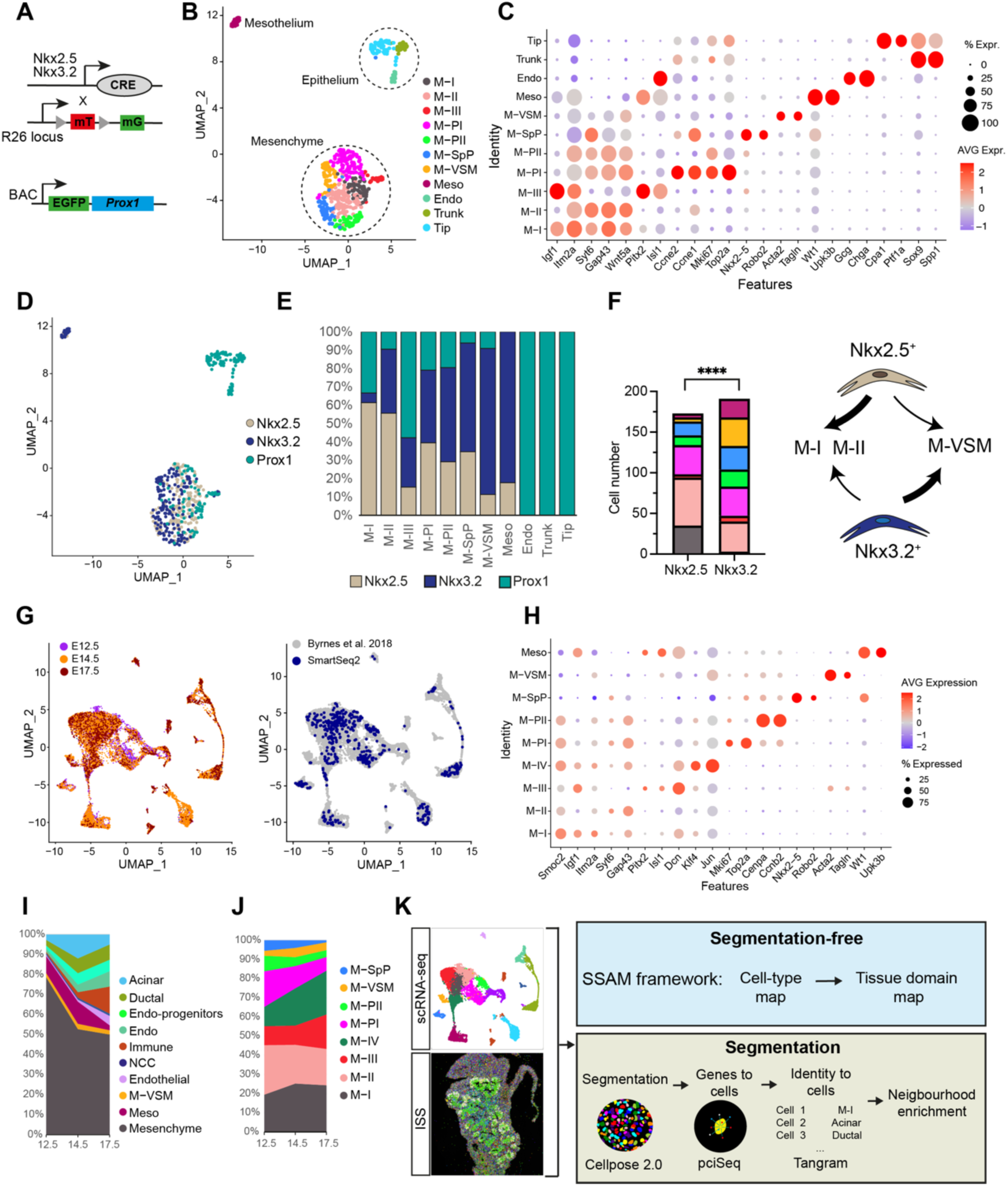
ScRNAseq analysis of Nkx2.5- and Nkx3.2-descendant mesenchymal cell types in the pancreas. (**A**) Smart-seq2 sequencing of GFP-positive cells FACS-sorted from E12.5 dorsal pancreas of *Nkx2.5-*Cre;*R26*mTmG, *Nkx3.2*-Cre;*R26*mTmG and Prox1-GFP BAC transgenic (Tg) mouse embryos. *Nkx2.5-*Cre and *Nkx3.2*-Cre recombined in specific subpopulations of the dorsal pancreas mesenchyme(*14,18*), whereas *Prox1* was expressed mainly in the pancreatic epithelium(*26,53*.) (**B**) UMAP plot showing clustering of Smart-seq2 scRNAseq profiles of embryonic pancreatic tissue at E12.5. After batch correction and clustering we identified 11 populations, including 3 epithelial (E) progenitor clusters (Tip, Trunk and Endocrine cells), 1 cluster expressing hallmark mesothelial genes (Meso), like *Upk3b* and *Wt1*, and 7 mesenchymal (M) clusters. Clusters are colour coded to indicate their annotated cell type. (**C**) Dot plot showing the expression level for selected marker genes for each cluster from (B). Proliferative clusters (M-PI and M-PII) are enriched in genes associated with cell cycle progression (*Mki67*, *Top2a*); the M-VSM cluster is enriched in genes expressed by mural cells (*Acta2*, *Tagln*); the spleno-pancreatic mesenchyme (M-SpP) cluster expresses high levels of transcription factors, such as *Nkx2.5* and *Tlx1*; M-I is characterized by high expression of *Itm2a* and *Igf1;* M-II is enriched in genes coding for collagens (*Col-1*, *Col-6*), non-canonical Wnt pathway (*Wnt5a*) and axon guidance (*Slit2*); M-III displays high levels of expression of *Pitx2* and *Isl1*, previously reported for a role in early stage pancreatic mesenchyme(*16,18*) (Table S1). Colour bar indicates the linearly scaled mean of expression level. (**D**) UMAP visualization of Smart-seq2 scRNAseq with cells coloured by the 3 Tg lines of origin. (**E**) Plot showing distribution of Tg cells across clusters (shown as %). Columns represent clusters; colours indicate the Tg line of origin. (**F**) Stacked bar plot showing the number of Nkx2.5- and Nkx3.2-mesenchymal cells contributing to each cluster (χ2, p<0,0001). Nkx2.5^+^ progenitors preferentially give rise to M-I and M-II clusters, while Nkx3.2^+^ progenitors to M-VSM cells. (**G**) UMAP plot of publicly available scRNAseq datasets(*16*) and Smart-seq2 of transgenic pancreatic rudiments (see Fig. 1B) after integration, coloured by embryonic stage (left panel) or by datasets (right panel). After batch correction and clustering, 17 cell populations were identified. These included known epithelial subtypes in the pancreas (acinar, ductal, endocrine progenitors and endocrine cells), neural crest-derived cells (*Phox2b*^+^), immune cells (*Cd74*^+^), endothelial cells (*Pecam1*^+^), blood cells (*Alas2*^+^), mesothelial (*Upk3b*, *Wt1*) and eight mesenchymal clusters, including M-I to M-III subtypes, two proliferative (M-PI and M-PII) clusters, one M-IV cluster (*Klf4*, *Jun*, *Fos*), one M-VSM cluster enriched in genes expressed by vascular mural cells (*Acta2*, *Tagln)* as well as pericytes, and one cluster expressing high levels of M-SpP genes. (**H**) Dot plot showing the expression level for selected marker genes for each mesenchymal (M) and mesothelial (Meso) clusters from the integrated scRNAseq dataset (as in Fig. 1B). Colour bar indicates the linearly scaled mean of expression level (table S2). (**I**) Proportion of cell clusters per developmental stage. (**J**) Proportion of mesenchymal cluster subtypes per developmental stage. (**K**) Schematics of the spatial transcriptomic analysis framework. HybISS and scRNAseq datasets were combined and analysed following segmentation-free and segmentation-based complementary approaches (See Materials & Methods section).

**Fig. S2.**
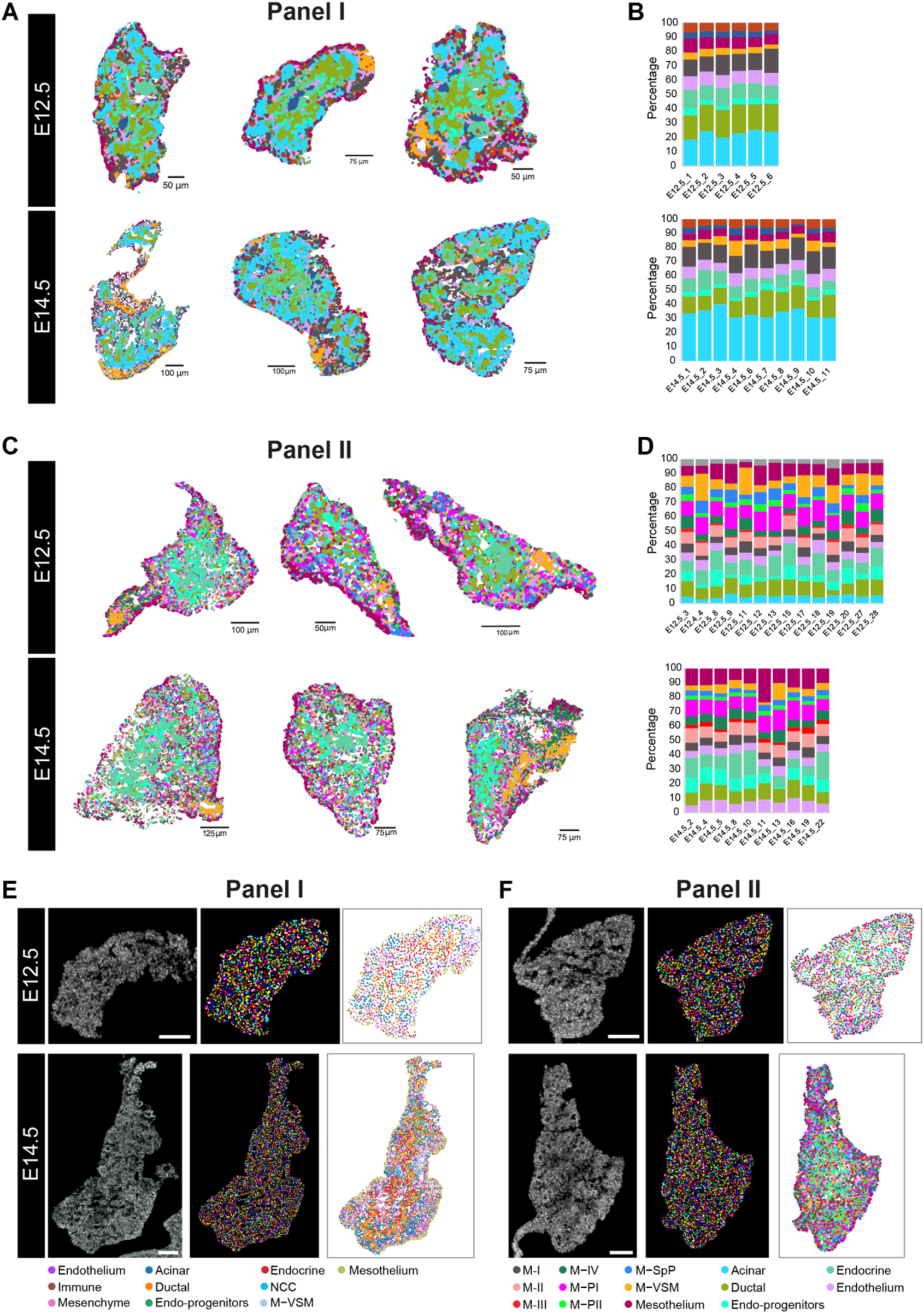
Segmentation-free and segmentation-based analyses of the HybISS datasets. (**A**) Representative SSAM-annotated cell maps of the HybISS Panel I experiment on E12.5 and E14.5 pancreatic tissue sections. (**B**) Cell type proportions per sample from E12.5 and E14.5 HybISS Panel I experiment. Colours represent single-cell clusters as shown below (E). (**C**) Representative SSAM-annotated maps of the HybISS Panel II experiments on E12.5 and E14.5 pancreatic tissue sections. (**D**) Cell type proportions per sample from E12.5 and E14.5 HybISS Panel II experiment. Colours represent single-cell clusters as shown below in (F). (**E,F**). Representative nuclear staining of E12.5 and E14.5 sections from Panel I (E) and II (F) (left). Nuclear segmentation staining generated using Cellpose 2.0 (middle); segmented nuclei are randomly coloured. Real position of annotated HybISS cells (right). Colours represent cell clusters as in Fig. 1A. Scale bar, 100μm.

**Fig. S3.**
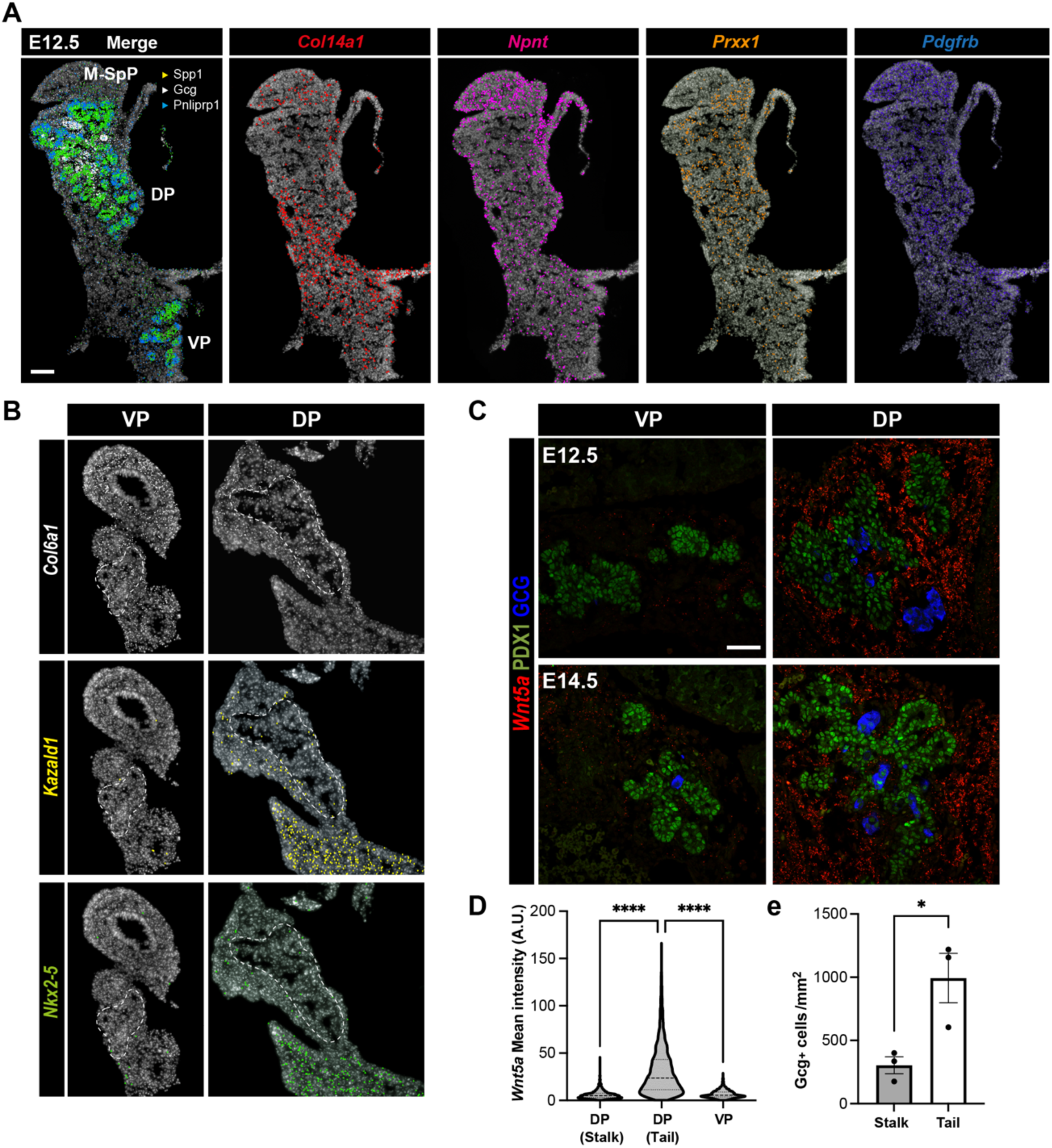
Dorsal and ventral pancreas show differences in their mesenchymal composition. (**A**) Representative HybISS images showing spatial distribution of indicated pancreatic reference genes (left panel) and mesenchymal marker genes (right panels) at E12.5. DP, dorsal pancreas; M-SpP, spleno-pancreatic mesenchyme; VP, ventral pancreas. Scale bar, 100μm. (**B**) Representative HybISS images showing spatial distribution of selected mesenchymal genes in VP (left) and DP (right) sections at E12.5. White dotted lines demarcate the pancreatic epithelium. (**C**) Representative confocal microscopy IF images of E12.5 and E14.5 DP and VP cryosections stained for indicated markers. Scale bar, 50μm. (**D**) Quantification of *Wnt5a mRNA* transcript levels on RNAScope-labelled DP at stalk and tail regions and VP tissues. AU, arbitrary units. n=3 embryos. ****p < 0.0001; One-way Anova test. (**E**) Quantification of glucagon (Gcg)^+^ cells in IF images. Cell counts were normalized to the average pancreatic epithelium area (mm2) and shown per area in stalk and tail regions of DP tissue. n=3 embryos. Error bars represent ± SEM. *p < 0.05; two-tailed unpaired t test.

**Fig. S4.**
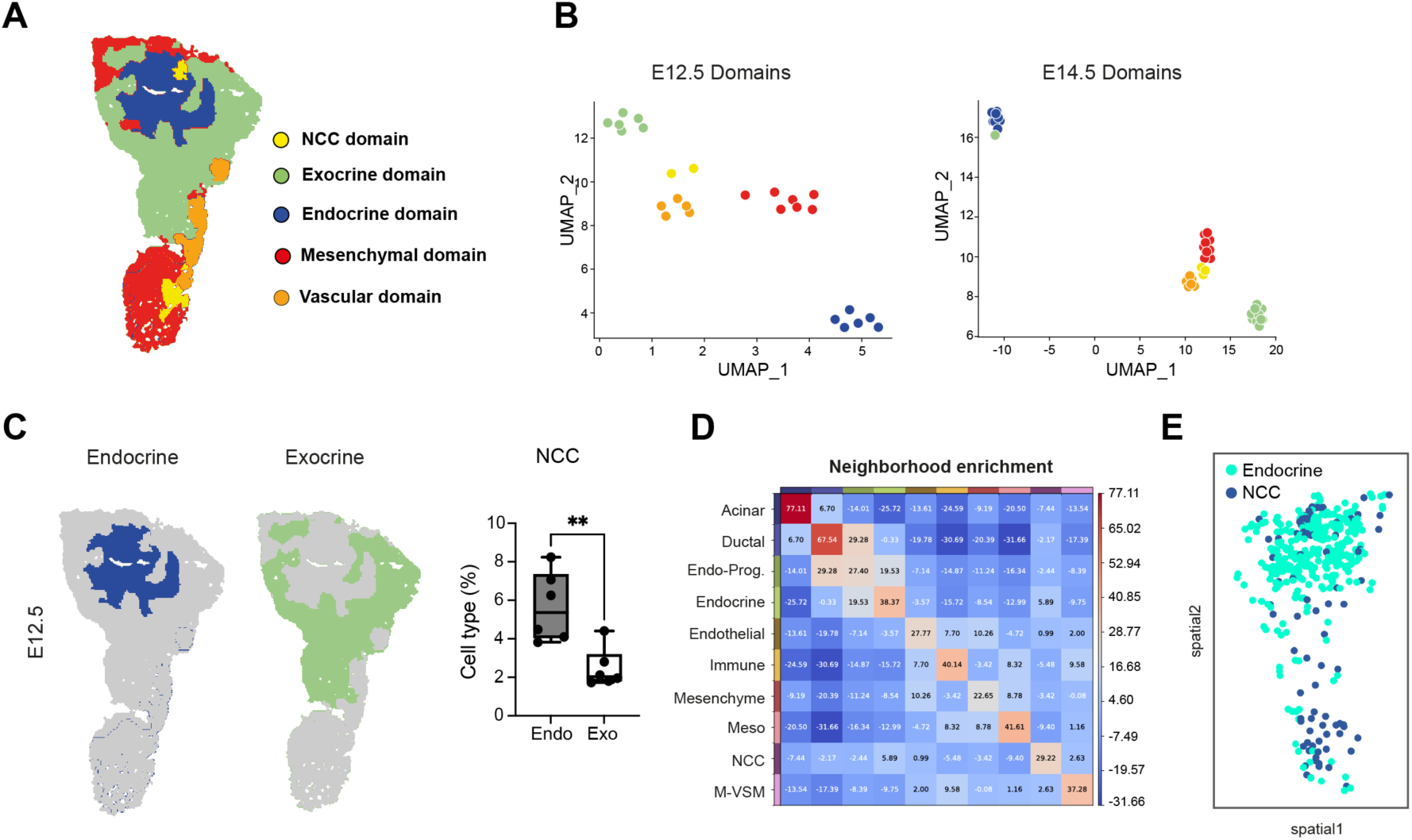
SSAM-based tissue domain annotation on images from Panel I dataset. (**A**) Representative SSAM-annotated tissue domain map from HybISS Panel I dataset at E12.5. (**B**) UMAP plots show annotated tissue domains at E12.5 and E14.5, identified by SSAM performed on all tissue images from Panel I dataset. Each point represents a predicted tissue domain from an individual tissue image. A list of tissue domains along with their cell composition is included in Table S4. (**C**) Representative endocrine and exocrine tissue domain maps of E12.5 pancreatic tissue (left) and quantification of NCC populations in each domain (right). **p < 0.01; two-tailed unpaired t-tests. (**D**) Neighbourhood enrichment score between cell types in E12.5 pancreatic tissue Panel I dataset (shown as mean; n = 6 tissue sections). Positive enrichment indicates proximity of a particular cell type to another one. Z-score indicates if a cluster pair is over-represented or over-depleted in the analysis. (**E**) Spatial distribution of NCC and endocrine cell types identified by HybISS on a E12.5 pancreatic tissue section. This is consistent with a previously reported role of NCC-derived cells in endocrinogenesis(*7*), highlighting their importance in the endocrine niche.

**Fig. S5.**
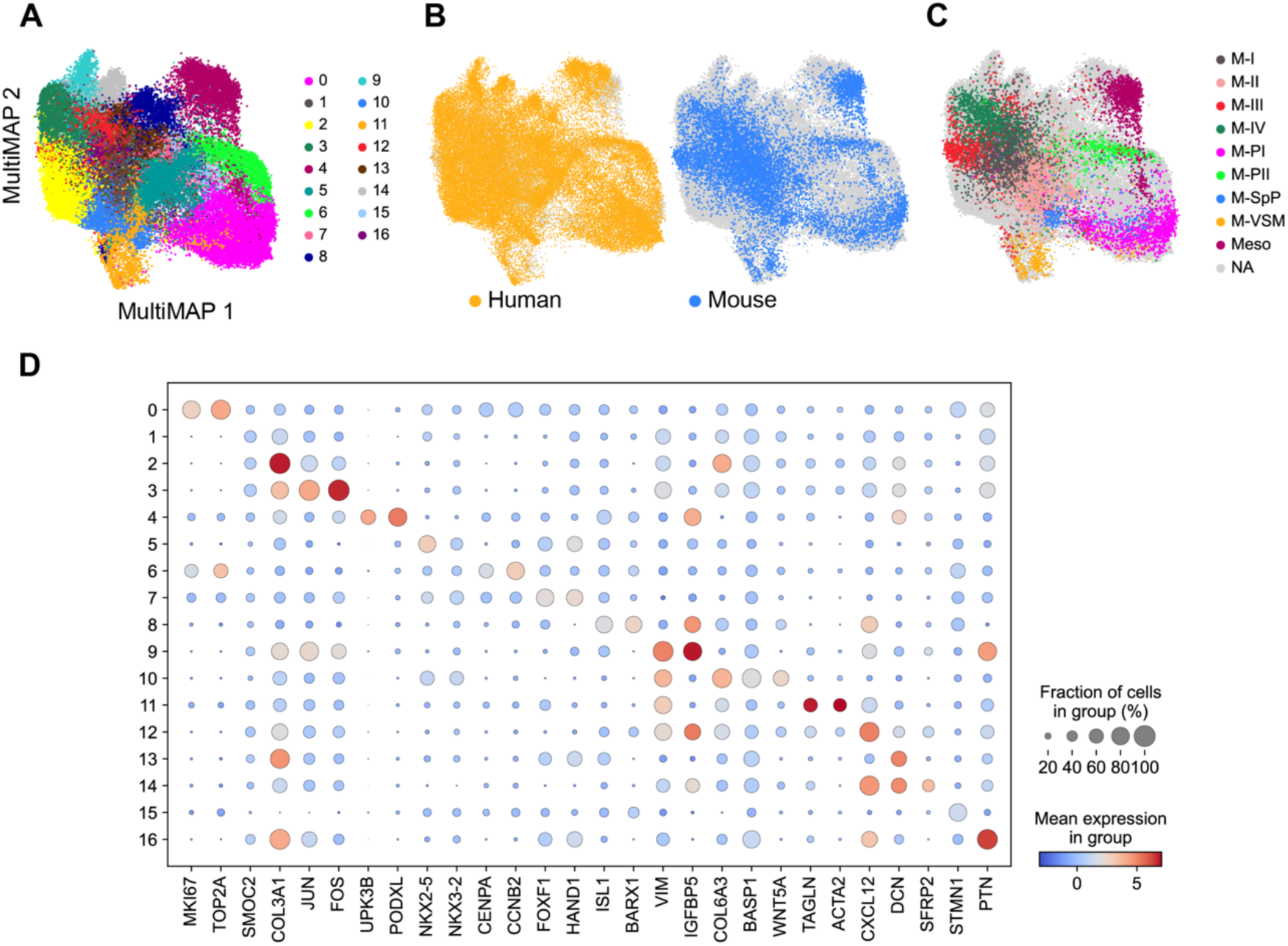
ScRNAseq comparison between human and mouse mesenchyme cells of the pancreas. (**A-C**) MultiMAP visualization of mesenchymal cells from human and mouse foetal pancreas coloured by cell cluster (A), species (B) or by annotated mouse clusters (C). N/A in (C) corresponds to human cells. (**D**) Dot plot showing the expression level of marker genes for each cell cluster from the integrated scRNAseq dataset [as shown in (A)]. Colour bar indicates the linearly scaled mean of expression level.

**Fig. S6.**
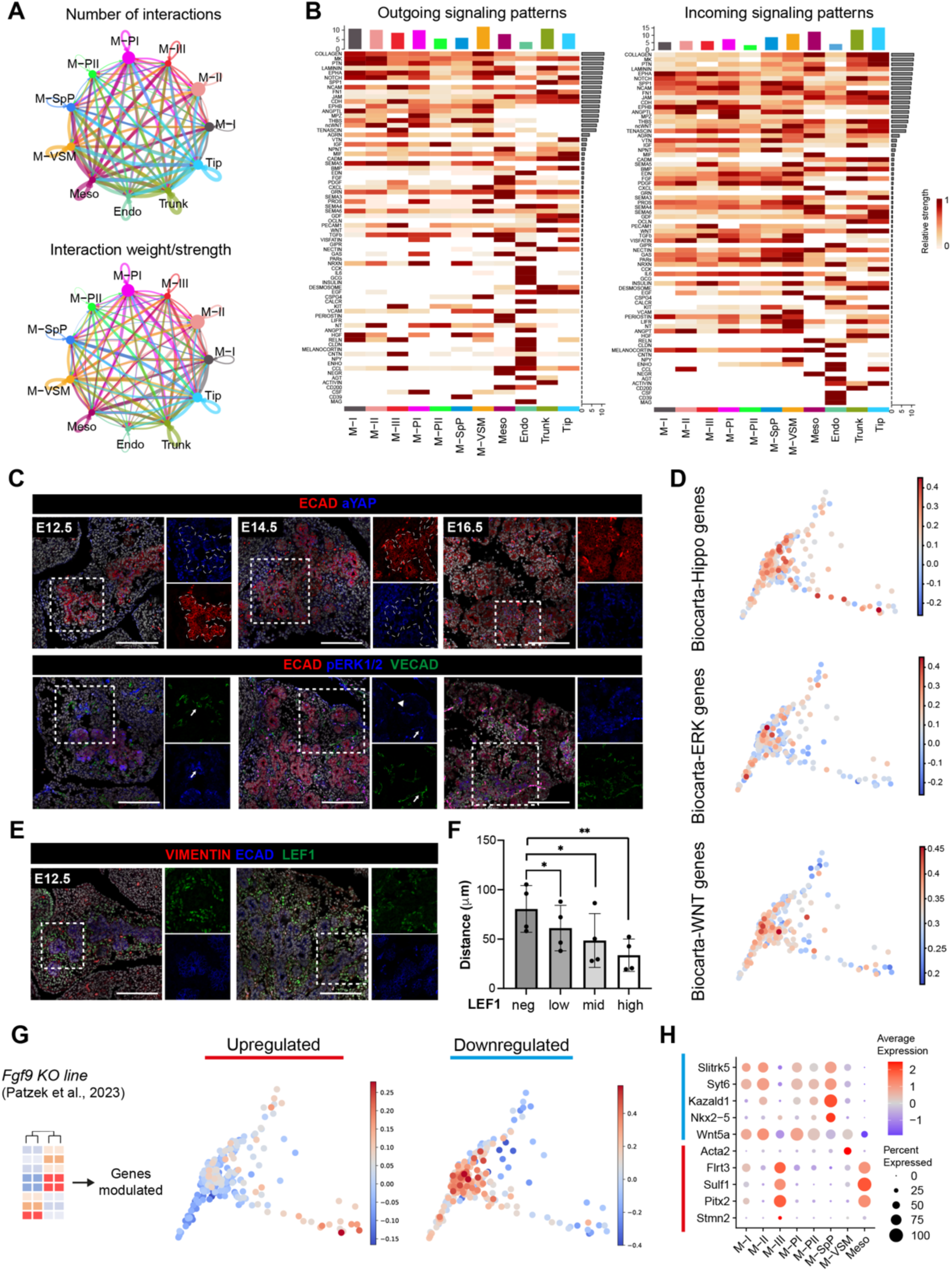
Cell signalling in the pancreatic mesenchyme. (**A**) Chord diagrams showing the cell-cell communication networks between pancreatic mesenchyme and epithelial cell types based on Cellchat analysis performed on our E12.5 Smart-seq2 dataset (see fig. S1A). (**B**) Heatmaps indicate the signalling strength of outgoing (ligands) and incoming (receptors) signalling pathways in each cell type. Signalling pathways are ordered by overall strength of signalling. (**C**) Representative confocal images of DP cryosections stained for the indicated signalling molecules. Right panels show boxed regions as single channels. White dotted lines demarcate the pancreatic epithelium. Arrows indicate p-ERK localization cells next to VECAD^+^ endothelium; arrowhead indicates p-ERK localization in mesothelial cells. Hoechst was used as nuclear counterstain. Scale bar, 150μm. (**D**) Diffusion plots of mesenchymal scRNAseq profiles showing average expression levels of normalized and log-transformed levels of gene modules from indicated signaling pathways based on BioCarta gene annotation (see Fig. 2E for M-clusters). (**E**-**F**) IF and quantification analysis of LEF1^+^ cell average distance to pancreatic epithelium (marked by ECAD staining). Scale bar, 150μm. LEF1^+^ cells were preferentially found next to the epithelium, suggesting a Wnt signaling gradient across the pancreatic mesenchyme. Right panels show boxed regions as single channels. Hoechst was used as nuclear counterstain. n=4 embryos. Error bars represent ± s.d.. *p < 0.05, **p < 0.01; One-way Anova test. (**G**) Gene signatures obtained from bulk RNAseq data of pancreatic tissue, isolated from *Fgf9* knockout (KO) mice(30), were projected into the Smart-seq2 E12.5 mesenchyme dataset diffusion plot. Genes downregulated in *Fgf9* KO were expressed in M-clusters closer to the spleen, whilst genes upregulated in the *Fgf9* KO were expressed in M-clusters closer to the duodenum, as defined by the transcriptional dynamics characterized in Fig. 2. (**H**) Dot plot showing a subset of the genes measured in (G) in E12.5 Smart-seq2 M-clusters. Colour bar indicates the linearly scaled mean of expression level.

**Fig. S7.**
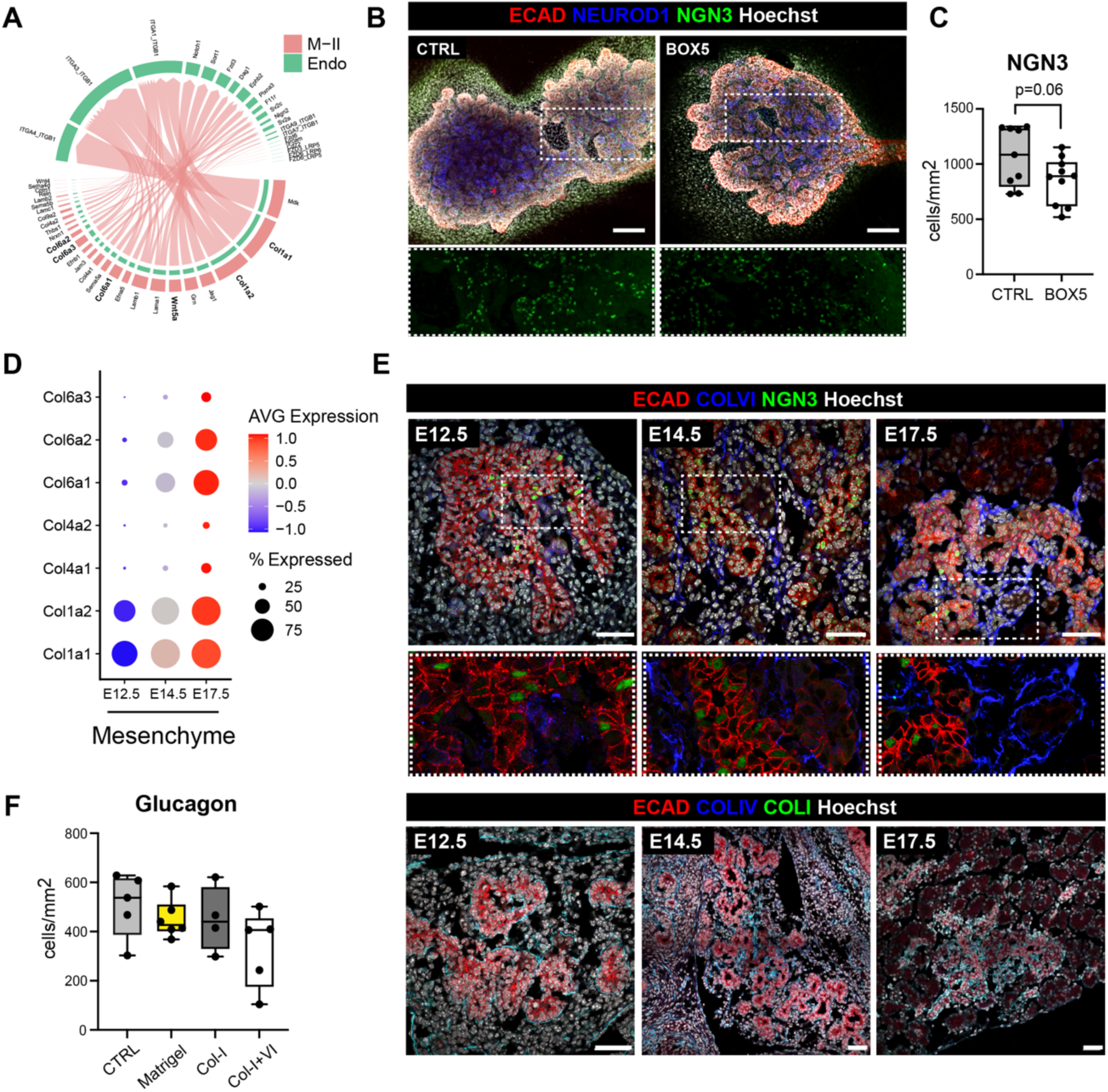
Characterization of M-II signaling niche. (**A**) Chord diagram showing putative cell-cell communication between pancreatic M-II and endocrine cell types based on E12.5 Smart-seq2 data. Endocrine cells are the ‘receiver’ cell type, expressing receptors and M-II are the ‘sender’ cells, expressing ligands. In bold, candidate molecules that were validated. (**B**) Whole-mount IF analysis for Neurogenin (NGN3), NEUROD1 and E-cadherin (ECAD) on mouse pancreatic explants treated for 2 days with BOX5 or left untreated as controls (Ctrl). Bottom panels show higher magnifications of the boxed region as NGN3 single channel (green). Hoechst was used as nuclear counterstain. Scale bars, 150μm. (**C**) Quantification of NGN3^+^ cells in pancreatic explants. Cell counts were normalized to the average ECAD^+^ epithelium area (mm2). n= 9-10 explants per condition. Two-tailed paired t-tests. (**D**) Dot plot showing the expression level of selected collagen genes in pancreatic mesenchymal cells per embryonic stage. Colour bar indicates the mean of expression level. (**E**) Representative confocal images of E12.5, E14.5 and E17.5 pancreatic cryosections immunostained for the indicated markers. Bottom panels show higher magnifications of the boxed region without Hoechst. An increase in collagen deposition was observed in the developing pancreas between E12.5 and E17.5, with Collagens I, IV and VI gradually accumulating around and inside the endocrine clusters. Scale bars, 50μm. (**F**) Quantification of Glucagon^+^ cells in pancreatic explants embedded in Matrigel, Collagen I, Collagens I and VI mix, or left unembedded. Cell counts were normalized to the average ECAD^+^ epithelium area (mm2). n=5-6 explants per condition. One-way Anova test.

## List of Supplementary Tables

**File name: Table S1**

Description: Differentially expressed genes of each cluster from the Smart-seq2 data set. Genes were calculated by comparing each cluster vs. all other clusters with Seurat’s FindAllMarkers() function.

**File name: Table S2**

Description: Differentially expressed genes of each cluster from the integrated embryonic data set. Genes were calculated by comparing each cluster vs. all other clusters with Seurat’s FindAllMarkers() function.

**File name: Table S3**

Description: Gene panels used for dRNA HybISS experiments. Each panel has its own tab.

**File name: Table S4**

Description: Tissue domain average cell type composition.

**File name: Table S5**

Description: Differentially expressed genes of each cluster from the adult mesenchyme data set. Genes were calculated by comparing each cluster vs. all other clusters with Seurat’s FindAllMarkers() function.

**File name: Table S6**

Description: Differentially expressed genes of each cluster from the adult-embryonic integrated mesenchyme data set. Genes were calculated by comparing each cluster vs. all other clusters with Seurat’s FindAllMarkers() function.

**File name: Table S7**

Description: List of primary and secondary antibodies, RNA probes, cytokines, media, and other reagents.

**File name: Table S8**

Description: Protocol, media and cytokines used for Beta Cell iPSC differentiation.

